# Organotypic endothelial adhesion molecules are key for *Trypanosoma brucei* tropism and virulence

**DOI:** 10.1101/2021.02.26.433042

**Authors:** Mariana De Niz, Daniela Bras, Mafalda Pedro, Ana Margarida Nascimento, Claudio A. Franco, Luisa M. Figueiredo

**Affiliations:** Instituto de Medicina Molecular Joao Lobo Antunes, Faculdade de Medicina, Universidade de Lisboa, Lisboa, Portugal; Departamento de Ciências da Vida, Faculdade de Ciências e Tecnologia, Universidade Nova de Lisboa, Lisboa, Portugal; Bioimaging Unit, Instituto de Medicina Molecular Joao Lobo Antunes, Faculdade de Medicina, Universidade de Lisboa, Lisboa, Portugal

**Keywords:** Tropism, vasculature, parasites, *Trypanosoma brucei*, intravital microscopy, endothelial cells

## Abstract

*Trypanosoma brucei* is responsible for lethal diseases in humans and cattle in Sub-Saharan Africa. These extracellular parasites extravasate from the blood circulation into several tissues. The importance of the vasculature in tissue tropism is poorly understood. Using intravital imaging and bioluminescence, we found that gonadal white adipose tissue and pancreas are the two main parasite reservoirs. We show that reservoir establishment happens before vascular permeability is compromised, suggesting that extravasation is an active mechanism. Blocking endothelial surface adhesion molecules (E-selectin, P-selectins, or ICAM2) significantly reduced extravascular parasite load in all organs and delayed host lethality. Remarkably, blocking CD36 had a specific effect on adipose tissue tropism that was sufficient to delay lethality, suggesting that establishment of the adipose tissue reservoir is necessary for parasite virulence. This works demonstrates the importance of the vasculature in a *T. brucei* infection and identifies organ-specific adhesion molecules as key players for tissue tropism.

## Introduction

Tissue-specific tropism within vertebrate hosts has been the focus of great interest in the field of parasitology in recent years. However, the cellular and molecular adaptations that allow parasite tropism are still poorly understood. For many parasites, tropism to specific organs is an essential step of their life cycle, as the organs provide a niche for persistence, latency or dormancy, massive replication and/or growth, protection from the host immune responses, or differentiation into alternative stages essential for completion of the life cycle, among others (reviewed in Boyett and Hsieh, 2014; Fernandes and Andrews, 2012; Guérin and Striepen, 2020; Lima and Lodoen, 2019; Onyilagha and Uzonna, 2019; Prudencio et al., 2006; Rénia and Goh, 2016; Silva Pereira et al., 2019; Venugopal et al., 2020).

*Trypanosoma brucei* is a parasitic organism transmitted by tsetse flies (*Glossina* spp.), responsible for human African trypanosomiasis (HAT) in humans, and nagana in other mammals. It requires two hosts to live and reproduce, namely, the insect vector, and the mammalian host (CDC, 2019). *T. brucei* invades the bloodstream and lymph, and disseminates across the host body (reviewed by Krüger and Engstler, 2018; Krüger et al., 2018). Several tissues and organs, including the adipose tissue (Trindade et al., 2016), skin (Alfituri et al., 2020; Capewell et al., 2016; Casas-Sánchez and Acosta-Serrano, 2016) and brain (Coles et al., 2015; Laperchia et al., 2016; Mogk et al., 2014; Myburgh et al., 2013), have been identified as important reservoirs of *T. brucei* due to the numbers of parasites harbored, the role they play in parasite transmission, or the associated pathology respectively (reviewed by Silva Pereira et al., 2019). In every instance, parasites live as extracellular forms in the parenchyma of those tissues. Yet, it is unclear the relative contribution of parasite tissue reservoirs to overall parasitemia and disease outcome.

Many parasites have evolved unique mechanisms to colonize tissues while avoiding being eliminated by the spleen. The relevance of the vascular endothelium for tropism and parasite dissemination has been studied in detail in the context of other parasites. For instance, *Plasmodium*-infected red blood cells (RBCs) undergo sequestration in the vasculature of several organs as a means to avoid spleen-mediated elimination (reviwed in Engwerda et al., 2005; del Portillo et al., 2012). Multiple endothelial receptors, including ICAM1, EPCR, PECAM1, CSA and CD36, have been well characterized and associated with sequestration in specific tissues in both human and rodent malaria infections (Fonager et al., 2012; Hviid and Jensen, 2015; De Niz et al., 2016; Smith et al., 2001). While *Plasmodium* asexual blood stages are, as far as we know, unable to cross the vascular endothelium, parasites such as *Toxoplasma gondii* invade endothelial cells and replicate within them as a step preceding dissemination into the extravascular niche of target organs, where parasites differentiate into quiescent forms (Konradt et al., 2016). *T. brucei* extravasation has been studied in the brain using *in vitro* models. Two key findings include that a trypanosome-derived cathepsin L-like cysteine protease (brucipain) is required for traversal of the brain-blood barrier; and that vascular permeability does not correlate with parasite dissemination in the brain (reviewed in (Kristensson et al., 2010)).

Here, we used intravital imagining to study the importance of the vascular endothelium in the establishment of *T. brucei* tissue reservoirs *in vivo*. Our data provide an unprecedented overall view of the dynamics of host-parasite interactions with the vascular endothelium and its relationship to parasite tropism. We identified the pancreas as a new reservoir for *T. brucei*, which, together with the gonadal white adipose (g-WAT) tissue, constitutes the organs with highest parasite load. We showed that g-WAT and pancreatic tropism is independent of organ-specific vascular density or vascular permeability. Instead, we found that *T. brucei* extravasation depends on several adhesion molecules expressed at the surface of endothelial cells. Finally, we identified CD36 as an adhesion molecules that specifically affected extravasation into the adipose tissue, demonstrating for the first time that the establishment of adipose tissue reservoir is a key virulence mechanism with great impact in disease outcome. Our analysis demonstrates for the first time that the vasculature plays an essential role in *T. brucei* infection and that the establishment of tissue reservoirs increases the fitness of parasites and disease severity.

## Results

### Gonadal white adipose tissue and pancreas are largest reservoirs of extravascular parasites

Several organs have been linked to T. brucei tropism (Silva Pereira et al., 2019); however, little is known about the dynamics of reservoir establishment. To evaluate parasite distribution in mice with high temporal resolution, we used a triple reporter parasite line that simultaneously expresses TY1, TdTomato and Firefly luciferase (Calvo-Alvarez et al., 2018). We injected 2×10^3^ parasites intraperitoneally in each mouse, and we followed the parasitemia *in vivo* in C57BL/6 mice and C57BL/6 Albino mice, the latter being a more suitable model to measure bioluminescence (Curtis et al., 2011; Doyle et al., 2004) (**Figure S1A left panel**). In both mouse strains, we observed high parasitemia at days 5-8, and then later in infection, as previously reported (Calvo-Alvarez et al., 2018). Both strains of mice showed similar survival times, with 50% survival being reached by day 22-23 post-infection (**Figure S1A middle panel**). Therefore, for all experiments discussed in this paper, we limited the infection to day 20.

In parallel to parasitemia, we followed up *T. brucei* infection by whole-body luminescence imaging in C57BL/6 Albino mice (**Figure 1A**). The relative whole-body luminescence correlated very well with blood parasitemia (R^2^= 0.82, p < 0.001) (**Figure S1A right panel**). To determine relative parasite load in individual organs, we dissected each organ following injection of luciferin, and performed *ex vivo* luminescence imaging (**Figure 1B, Figure 1C, Figure S1B**). Three white and two brown adipose tissues (WAT and BAT, respectively) were collected (schematic shown in **Figure S1C**). Animals were not perfused prior to organ excision, therefore the signal measured includes both intra- and extravascular parasites (**Figure 1C, Figure S1B**). Gonadal white adipose tissues (g-WAT), pancreas, and lungs (marked with arrows, **Figure 1C**) were the organs that presented the highest bioluminescence values, suggesting they contain on average more parasites than other organs throughout infection (**Figure 1C, Figure S1B**).

**Figure 1.**
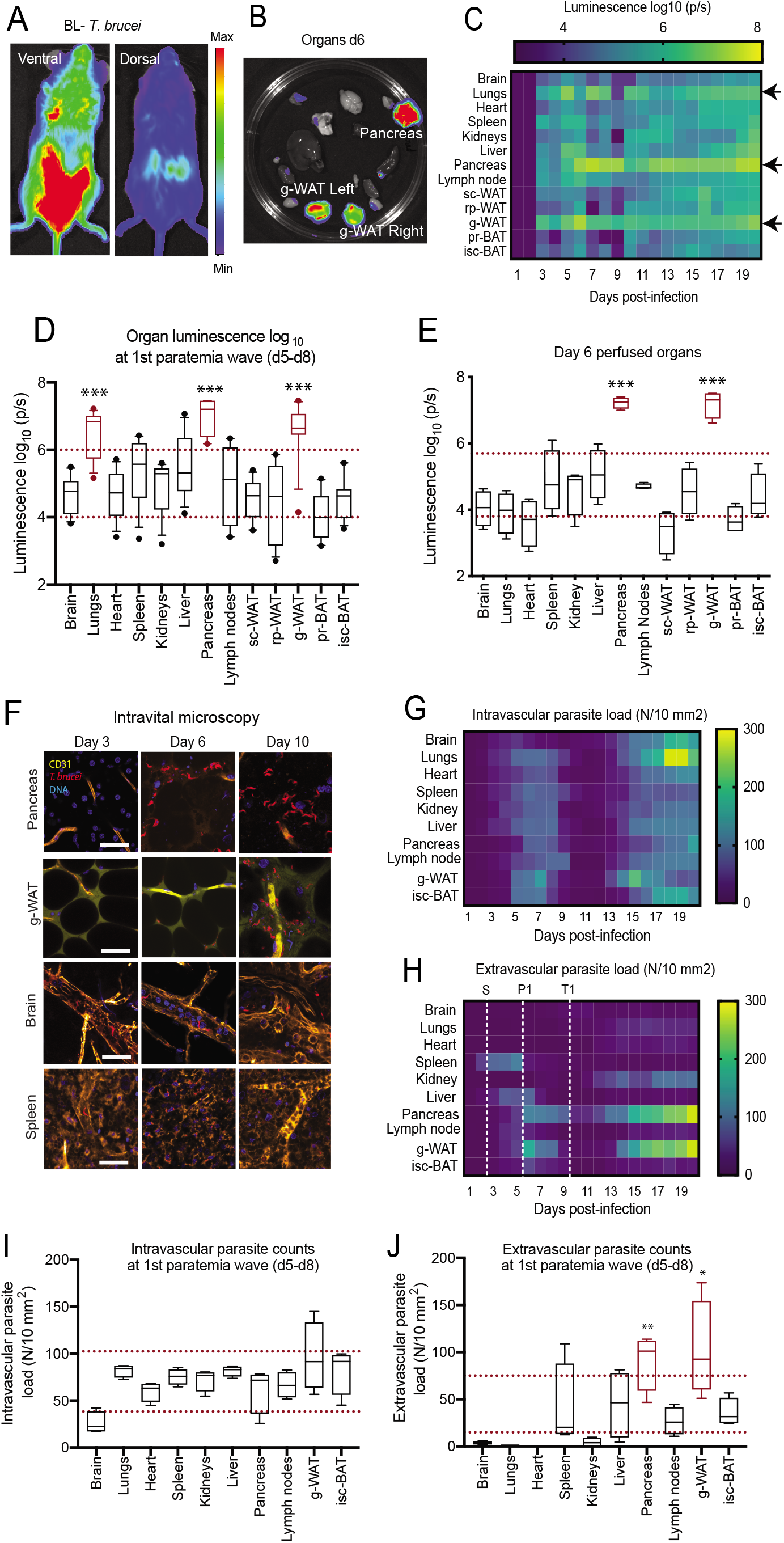
Identification of two *Trypanosoma brucei* extravascular reservoirs by bioluminescence and intravital microscopy. **(A)** Representative image of whole body bioluminescence in ventral and dorsal positions (left) using an IVIS Lumina system. **(B)** Organs were collected from infected mice (day 6 post-infection) previously injected with RediJect luciferin. **(C)** Bioluminescence of individual organs expressed as log_10_ light units normalized per tissue area. P7S: photons per second. The detected signal is a mixture of extravascular and intravascular parasite loads. **(D)** Box-plots of combined light unit measurements during the first wave of parasitemia (days 5 to 8). Red dotted lines show the interquartile range considering all organs. Organs above the upper threshold are considered significantly enriched (lungs, pancreas and g-WAT). **(E)** Bioluminescence of individual organs obtained from mice that were perfused at day 6 post-infection. **(F)** Immunofluorescence representative images obtained from intravital live imaging of parasites (TdTomato reporter, red) in g-WAT, pancreas, brain and spleen at 3 different days of infection. Vessels are labeled with anti-CD31 antibody (yellow), interstitial space with FITC-Dextran and nuclei with Hoechst (blue). **(G-H)** Heat-map of parasite density, calculated by confocal microscopy as the number of detected parasites per area inside (F) or outside vessels (G). Three dotted white lines are used to denote three key events during infection namely start of extravascularization (S, day 2), first peak of infection (P1, day 6) and the beginning of the reduction of intravascular parasite density (trough) (T1, day 10). **(I-J)** Box-plots of combined quantifications of *T. brucei* per area of tissue inside (H) or outside (I) vessels during the first wave of parasitemia (days 5 to 8). Red dotted lines show the interquartile range considering all organs. Organs above the upper threshold are considered significantly enriched (none for intravascular parasites; pancreas and g-WAT for extravascular parasites). For all graphs, significance is shown as p < 0.001 (***), p < 0.01 (**), p < 0.05 (*). Scale bar is 50 μm.

For quantitative comparison, we show pooled luminescence values of all organs during the first peak of parasitemia (days 5-8 post-infection) (**Figure 1D**). The lungs, pancreas and g-WAT showed an overall average of 10^6^-10^8^ luminescence units, around 30-fold higher than the average of other organs (which had an accumulated average of 10^4^ – 10^6^ luminescence units, **Figure 1D**). A strong accumulation of parasites was detected in these, and most other organs, since day 3 postinfection (**Figure 1C, Figure S1B**). Without perfusion, among adipose tissues, the most striking observation was that throughout infection the parasite signal in the g-WAT was around 60-250-fold higher than other WAT depots (p < 0.001 for all comparisons to g-WAT). Between the two remaining WAT depots (sub-cutaneous and retroperitoneal), the luminescence values were not significantly different (paired t-test, p = 0.82). Between the two BAT depots (peri-renal and interscapular), the luminescence values were also not significantly different (paired t-test, p = 0.56). Global comparisons between WAT and BAT depots showed that luminescence in BAT depots was significantly lower than WAT depots (p = 0.04) (**Figure S1C**). The pancreas was the organ with the highest accumulated daily intensity, being on average 2.8-fold higher than g-WAT and 22-fold higher than lungs considering the 20 days of infection.

In the bloodstream, parasites are at very high concentrations (**Figure S1A**). To discriminate the bioluminescent signal between intravascular and extravascular compartments, we perfused mice prior to measuring bioluminescence to remove all signal emanating from intravascular parasites on day 6 post-infection (**Figure 1E**). In perfused mice, only g-WAT and pancreas showed significantly high luminescence. Upon perfusion, g-WAT was 200-4800-fold higher than the four other WAT or BAT depots (p < 0.001 for all comparisons to g-WAT). Upon perfusion, at day 6 post-infection the pancreas was equally enriched as the g-WAT (p=0.25), and 1000-fold higher than lungs (p<0.001). Notably, while in non-perfused mice the lungs show very high bioluminescence values (**Figure 1D**), the signal intensity decreased significantly after perfusion by 200-fold (p < 0.001) (**Figure 1E**), suggesting that most parasites in the lungs remained intravascular. In conclusion, our data show the g-WAT and pancreas are the largest extravascular parasite reservoirs.

To confirm the bioluminescence results, we determined the intra- and extravascular location of parasites in each organ at single cell level, using intravital microscopy and *ex vivo* confocal microscopy (**Figure 1F-1J**). Prior to each imaging session, we injected mice with the pan-vascular marker CD31 conjugated to A647 (orange in the images), and/or FITC-Dextran to be able to define the intra- and extra-vascular spaces (selected organs are shown in **Figure 1F**). While intravital microscopy allows quantifications at single cell resolution, the penetration depth achieved limits the total area of tissue that can be covered, therefore explaining to some extent, differences observed with bioluminescence. Conversely, while bioluminescence allows imaging the full organ, it lacks the single cell resolution that intravital microscopy allowed. Taking this into consideration, using intravital microscopy, intravascular parasite density (parasites/mm^2^) showed a similar pattern of parasitemia in all organs, with two parasitemia “waves” in peripheral blood (**Figure 1G**), in accordance to measurements performed by haemocytometer (**Figure 1C, Figure S1A**). Quantitative values corresponding to intra- and extra-vascular parasites during the first parasitemia wave (day 5-day 8) are shown in **Figure 1I** and **IJ**, respectively, confirming significant enrichment in the g-WAT and pancreas. Throughout infection, in total, the number of parasites per mm^2^ found in the extravascular space of g-WAT and pancreas corresponds to around 40% of the sum of extravascular parasites counted per mm^2^ in the 13 organs, while the three white adipose tissue depots together contribute 48%. Supplementary videos 1-2 show *T. brucei* in intravascular and/or extravascular locations.

The intravital-imaging analysis identified 4 different phenotypes of intravascular and extravascular *T. brucei* distribution across organs (**Figure 1G-1J, Figure S2**). The two organs we identified as reservoirs (pancreas and g-WAT), together with the sub-cutaneous and the retroperitoneal WAT (**Figure S2A**), form group 1, in which there is proportionally more parasites in the extravascular space (blue) than intravascularly (red) throughout most of infection. The single cell measurements are consistent with the bioluminescence assays performed with and without perfusion (**Figure 1E**). Group 2 (including the two brown adipose tissue depots) (**Figure S2B**) has a pattern similar to group 1, in which the intra- and extravascular compartments present two waves of parasite populations with a minimum around day 11. In contrast to group 1, group 2 has more parasites in the intravascular space than in the extravascular space. Given the consistency of patterns between the BAT and WAT groups, henceforth, we only use one representative of each tissue category (gonadal for WAT, and interscapular for BAT). In a third group (brain, lungs, heart, and kidneys) (**Figure S2C**), the vast majority of parasites reside in vessels, with a small population colonizing the extravascular space mostly at later times post-infection after the first peak of blood parasitemia. Finally, the last group of organs (spleen, lymph nodes, and liver) (**Figure S2D**) showed an early extravascularization of parasites prior to the peak in the peripheral blood, but little enrichment later in infection. This enrichment preceded the first peak of peripheral blood parasitemia. Importantly, following this early peak, all the organs belonging to this group underwent significant remodeling, including enlargement (lymph nodes and spleen), appearance of white foci (liver and spleen), and increased vascular permeability (liver, lymph nodes and spleen – discussed in the next section).

Altogether, these results indicate that parasite distribution in intra- and extra-vascular spaces is very heterogenous across organs and it varies with time. Importantly, in terms of establishment of an early tissue reservoir, we conclude that g-WAT and pancreas parasite reservoirs are established since day 3 post-infection and that both remain the most parasitized organs throughout infection.

### *T. brucei* enrichment in the pancreas and white adipose tissues is not correlated to vascular density

Next, we investigated the reasons for g-WAT and pancreatic tropism. First, we explored if there were differences in vascular density between organs. If the establishment of parasite reservoirs depends on the probability that parasites have of interacting with and crossing blood vessels, we postulated that extravascular parasite density would correlate with vascular density. To test this hypothesis, we performed a combination of intravital microscopy and *ex vivo* imaging in tissue sections of all organs throughout infection. Vascular density was measured by determining the total tissue area imaged, and the percentage of that area that was constituted by vessels (marked by CD31, an endothelial receptor, and FITC-Dextran, a large polysaccharide that stays inside vessels unless there is increased vascular permeability) (**Figure 2A**).

**Figure 2.**
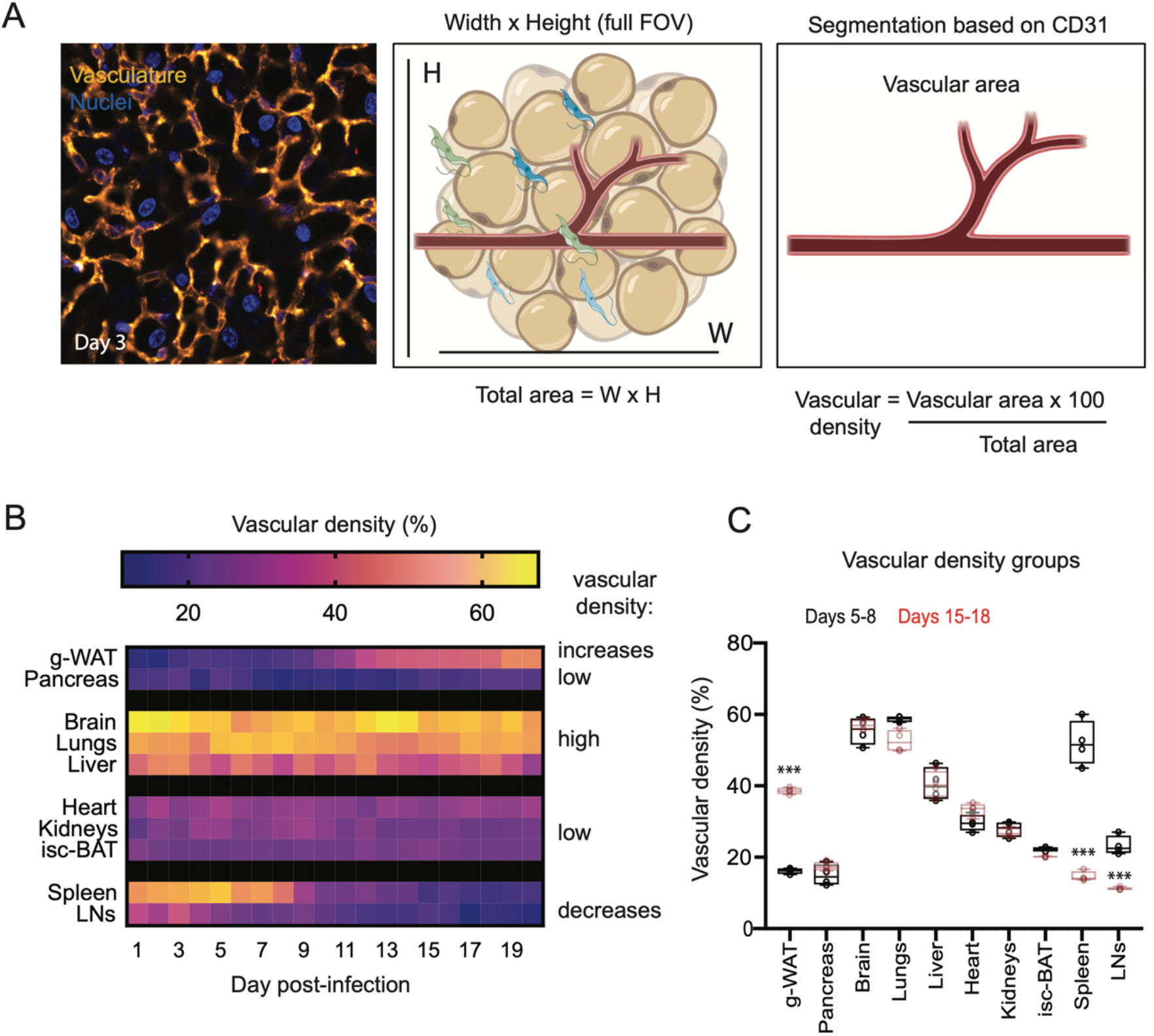
Changes in organ vascular density during *T. brucei* infection. **(A)** Vasculature is labelled with an antibody anti-CD31 (yellow), while nuclei are labelled by Hoechst (blue). Vascular density is calculated as the area occupied by the vessel divided by the total area of the field of view were performed following vascular segmentation based on the marker CD31 (left panel) as shown in the schematic (middle and right panels). **(B)** Heat-map of vascular density (calculated as defined in (A) throughout 20 days of infection. Organs are divided into groups, namely a) increasing vascular density, high (>35%), low (<35%), and decreasing vascular density. **(C)** Box plot showing quantitative values corresponding to (B) for days 5-8 (black) and 15-18 (red), showing the increase and decrease in vascular density in groups 1 and 4. For all graphs, significance is shown as p < 0.001 (***), p < 0.01 (**), p <0.05 (*).

In non-infected animals and consistent with literature (Cook, 1965), we found that organs such as brain, lungs, liver, spleen and lymph nodes are highly vascularized (30-60% of their area consists of vessels), while adipose tissue depots, pancreas, heart and kidneys are less vascularized (12-30%) (**Figure 2B-2C, Figure S3**). **Figure 2C** shows quantitative values for vascular density during the first parasitemia wave (days 5-8, black boxes) and the last parasitemia wave (days 15-18, red boxes). During infection, vascular density remained unchanged in most organs, including pancreas. In contrast, vascular density in g-WAT significantly increased from 13% at day 1 to 49% at day 20 post-infection (**Figure 2B, Figure S3**) and it significantly decreased in spleen and lymph nodes from 51 and 36% respectively at day 1, to 16 and 8% respectively at day 20 post-infection (**Figure 2B, Figure S3**). The increase in vascular density observed in the g-WAT coincides with (and might contribute to) tissue shrinkage by up to 82% probably due to extensive lipolysis observed as infection progresses (also observed in other WATs) (Trindade et al., 2016). Conversely, the decrease in vascular density observed in the spleen and lymph nodes could be due to the dramatic tissue enlargement (increase of 60% and 210%) that these organs undergo as infection progresses.

When we compared the variation over time of the organ vascular density with the organ extravascular parasite load during 20 days of infection (**Figure 1H**), we found a significant correlation for some organs (g-WAT, pancreas, kidneys, isc-BAT, and spleen) but not others (heart, brain, lungs, liver and lymph nodes). However, when we considered whether vascular density correlated with reservoir establishment (i.e. considering extravascular parasite load only the first 6 days of infection), we found only a significant correlation for the spleen (R^2^ = 0.92, p <0.01). Conversely, considering only these days, the g-WAT (R^2^ =0.13, p = 0.54) and pancreas (R^2^ = 0.006, p = 0.9) showed no strong correlation. Altogether, we can conclude that vascular density is not the reason why white adipose tissues and pancreas are the main reservoirs early in infection.

### Parasite distribution across blood vessels is heterogeneous

Capillaries and post-capillary venules of several organs have been shown as important locations for *Plasmodium* sequestration (overview by Franke-Fayard et al., 2010). Thus, we asked whether the type of vessel allowing parasite traversal is important for *T. brucei* tropism. In mammals, the vasculature is comprised of vessels of different diameters, of either arterial or venous origin (**Figure 3A**) (OpenStax; Tucker WD, Arora Y, 2020). We hypothesized that in the adipose tissue and pancreas, parasites may accumulate in a specific type of vessel, which could favor crossing from the blood into the extravascular space. To address this question, we injected uninfected and infected mice with exogenously-labeled RBCs (as the basis for normalization for quantifications), and monitored their distribution across all vessel types, which were marked either with the pan-endothelial marker CD31, the arterial marker Ephrin B2 or the venous marker EphB4, conjugated to A647 (**Figure 3B**). It has been shown that the distribution of RBCs is proportional to the caliber and hemodynamics in each vessel (Carlson et al., 2008; Chien, 1970; Chien et al., 1966; Coppola and Caro, 2009; Dobrin, 2011; Secomb, 2016), which are also the conditions faced by intravascular parasites. For each day of the infection and for each organ, we surgically stopped blood flow and we quantified the number of parasites relative to the number of labelled RBCs. Vessels were classified according to their caliber either considering the total vasculature (based on marker CD31, **Figure S4**) or by venous and arterial identity (based on markers EphB4 and Ephrin B2, **Figure 3E-3H**): small (0-10 μm) comprising capillaries, medium (10-20 μm) corresponding to arterioles and venules, and large (>20 μm) comprising arteries and veins. The longitudinal results of parasite concentration in each type of vessel are displayed as heat-maps, while the overall distribution for each condition is represented by violins plots (**Figure 3, Figure S4, Figure S5**).

**Figure 3.**
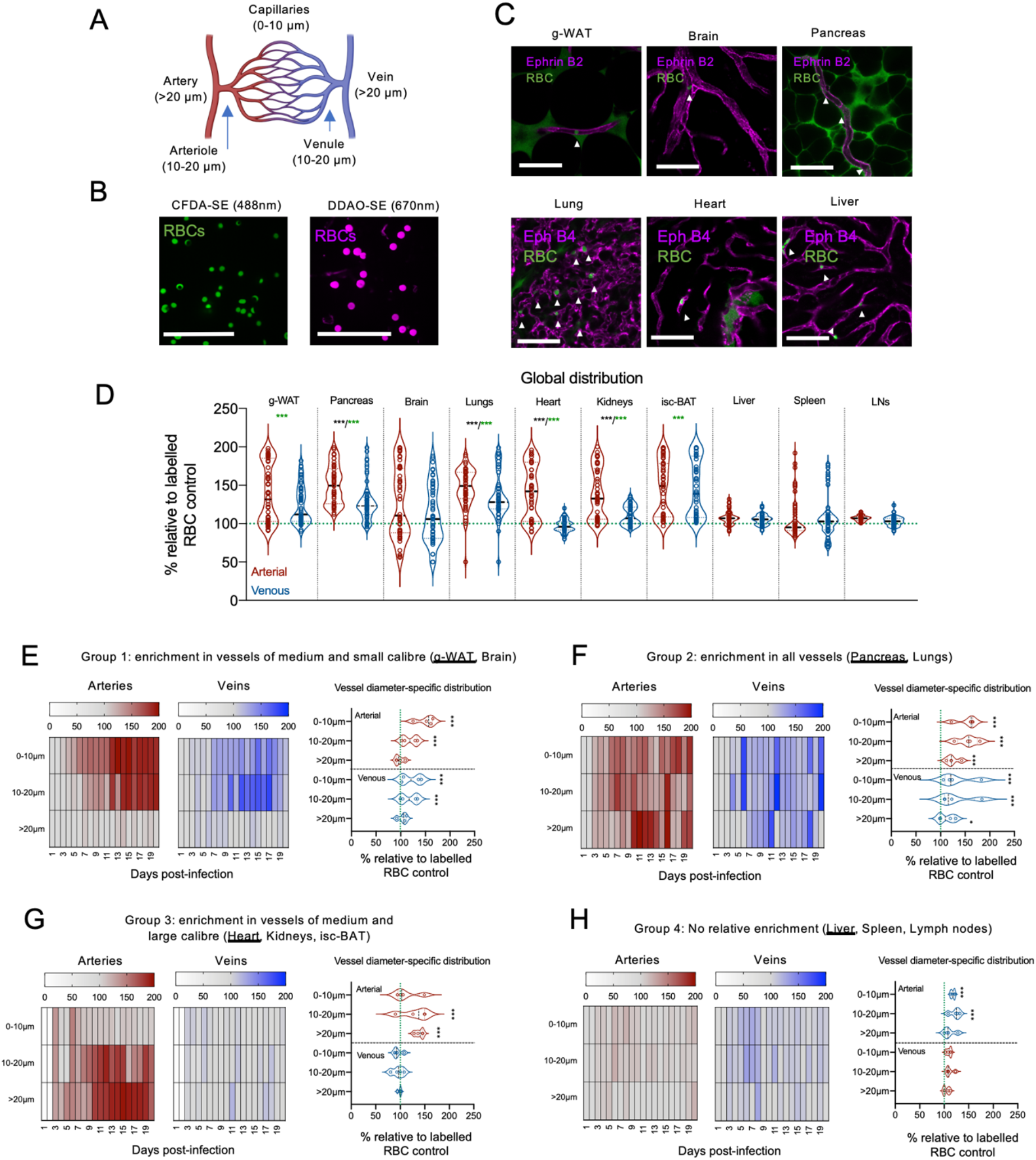
Distribution of *T. brucei* across arterial and venous vasculature is organ-specific. **(A)** Schematic of arterial and venous vasculature, their deriving branches and respective diameters. **(B)** Representative microscopy images of CFDA-SE and DDAO-SE-labeled RBCs, which were injected into mice to allow the measurement of the relative concentration of parasites in a given vessel. Parasite concentration was calculated as the number of parasites per RBC. **(C)** Representative immunofluorescence images show arteries as Ephrin B2-stained vessels, while veins are Ephrin B4-stained vessels. CFDA-SE-labeled RBCs are shown in green (marked by arrows). **(D)** Violin plots showing quantifications of global parasite concentration relative to RBC in arterial or venous vasculature for the 10 organs analysed in this study (green dotted line indicates that there are 100 parasites per 100 RBC). Significance relative to RBC normalizers is shown as green asterisks. Significant differences between arterial or venous groups is shown as black asterisks. Significance is defined as follows: p < 0.001 (***), p < 0.01 (**), p < 0.05 (*).

**(E-H)** Heat-maps of parasite distribution in vasculature, relative to labeled RBCs. Panels are divided in arterial and venous vasculature. Violin plots show parasite concentration relative to RBC according to vessel diameter (0-10, 10-20 and >20 μm) and vessel type (arterial or venous). Organs were grouped by similar parasite distribution defined in Figure S4. Significance between vasculature of different diameters per vessel type is defined as follows: p < 0.001 (***), p < 0.01 (**), p < 0.05 (*).

Vessel caliber was the most important variable that contributed to heterogenous parasite distribution in the vasculature, when looking at total vasculature. We identified 4 phenotypes of parasite enrichment across organs (shown in **Figure S4**): group 1, enrichment in medium and small vessels (**Figure S4B**) in the brain and g-WAT; group 2, enrichment in all vessels (**Figure S4C**) in the pancreas and lungs; group 3, enrichment in medium and large vessels (**Figure S4D**) in the heart, kidneys and isc-BAT; and group 4, no relative enrichment (**Figure S4E**) in the liver, spleen and lymph nodes.

When taking into account arterial and venous identity, across the 20 days of infection, we saw that in g-WAT parasite concentration is similar in arteries and veins (p = 0.12) as is the case also for the brain (p = 0.16), lungs (p =0.14), isc-BAT (p = 0.93), liver (p = 0.68), spleen (p = 0.74) and lymph nodes (p = 0.74). In contrast in pancreas (p = 0.02), heart (p < 0.001), and kidneys (p = 0002) parasites were globally more concentrated in vessels of arterial origin (**Figure 3D**).

In g-WAT (**Figure 3E**), parasite concentration was highest in vessels of small and medium caliber of both arterial and venous origin, while in the brain the enrichment was mostly in arterial vasculature (**Figure S5A**). In the pancreas and lungs (**Figure 3F, Figure S5B**), parasites were highly enriched across arterial and venous vessels of all calibers. While the heart, kidney and isc-BAT (**Figure 3G, Figure S5C**) showed a higher parasite concentration in vessels of medium and large caliber, the heart mostly displayed enrichment in the arterial vasculature, while the kidneys and isc-BAT showed enrichment in both arterial and venous vasculature. Finally, in the spleen and lymph nodes (**Figure S5D**), parasite concentration was equivalent to the labelled RBC markers, showing no significant enrichment in any specific vessel type, while in the liver (**Figure 3H**) the arterial vasculature seemed to be enriched.

In conclusion, *in vivo* imaging of *T. brucei* in the vasculature of infected mice shows that the distribution of parasites across vessels is extremely heterogeneous, organ- and time-dependent. Our results are consistent with the notion that endothelial cells have unique transcriptomic signatures and features in each organ, a concept termed as organotypic vasculature (Augustin and Young Koh, 2017). It also further suggests that these specific features allow for differential distribution of parasites across organs. Yet, given that parasite distribution in g-WAT and pancreas is different between these two organs and it is not unique relative to other organs, it shows that reservoir establishment is not uniquely associated with parasite enrichment in a specific vessel type or caliber. In fact, organs with similar parasite distribution (i.e. g-WAT and brain, or the pancreas and lungs) do not form equivalent extravascular reservoirs. Thus, additional features should contribute to reservoir establishment.

### Extravascular reservoirs are established before vascular permeability is compromised

Vascular permeability is typically increased during an infection, and has previously been related to facilitating immune cell extravasation (Schnoor et al., 2015; Vestweber, 2015; Vestweber et al., 2014; Wessel et al., 2014). Yet, changes in vascular permeability have not been previously studied in a *T. brucei* infection. We hypothesized that increases in vascular permeability could regulate parasite extravasation preferentially in the g-WAT and pancreas, allowing tissue tropism. To investigate vascular permeability, we intravenously injected mice with CD31 and 70 kDa FITC-Dextran. Using both markers, we measured both the intravascular and extravascular mean fluorescence intensity (MFI) of FITC-Dextran using established methodology for intravital microscopy (Egawa et al., 2013). At basal conditions, albeit depending on each organ’s inherent vascular permeability, most of the FITC-Dextran will remain in the blood vasculature. If the organ’s vasculature becomes more permeable, FITC-Dextran will leak into the organ’s parenchyma (schematic representation shown in **Figure 4A**).

**Figure 4.**
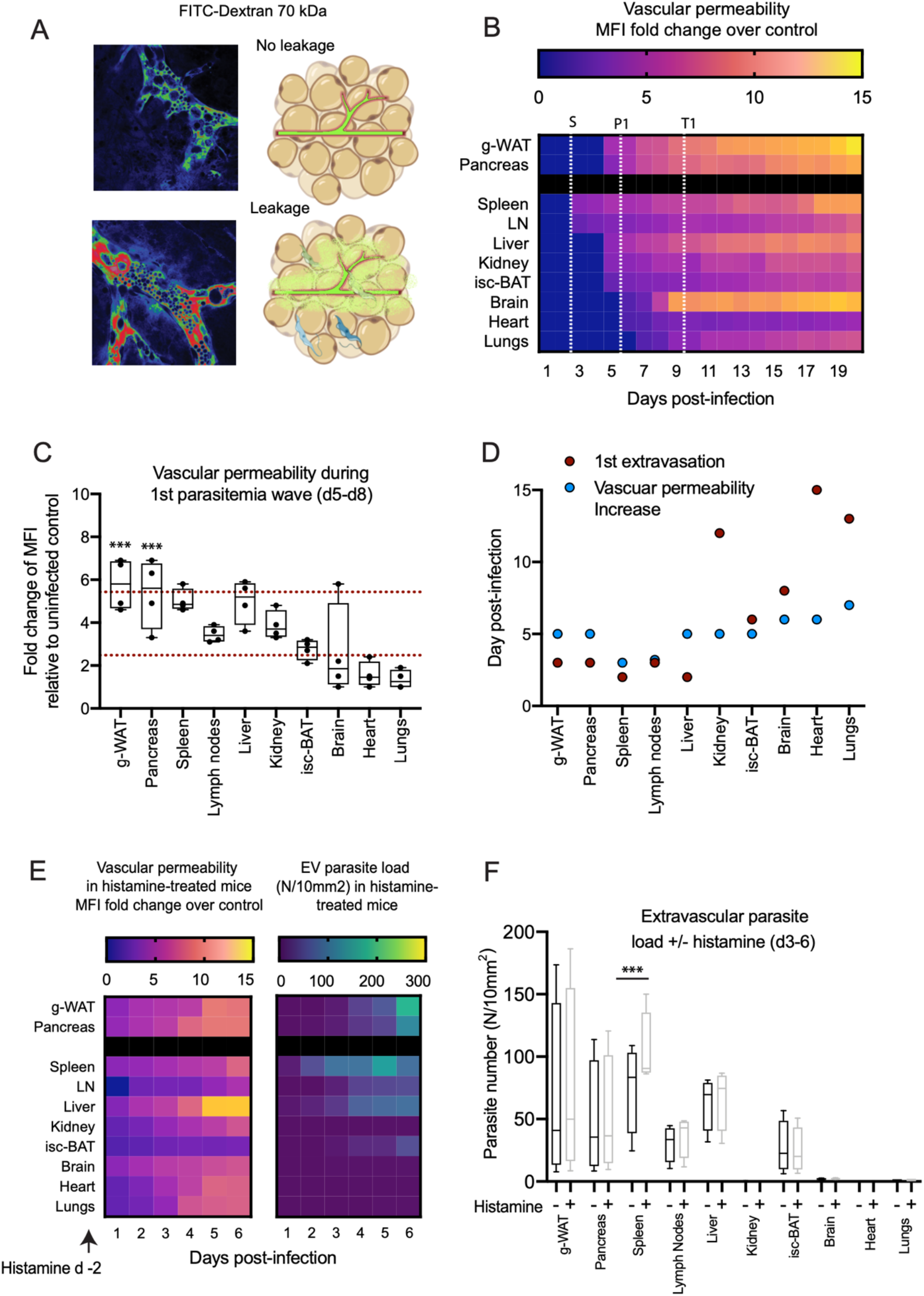
Changes in vascular permeability during *T. brucei* infection. **(A)** Vascular permeability was measured in every organ as an increase in the mean fluorescence intensity (MFI) emitted by the 70kDa FITC-Dextran in the extravascular space. **(B)** Heat-map of vascular permeability every day of infection. Vascular permeability is calculated as the FITC-Dextran MFI in the extravascular space at each day post-infection relative to FITC-Dextran MFI in uninfected control (baseline values are shown in Figure S4). The white dotted lines show the markers defined in Figure 1G for the start of reservoir establishments (S, day 3), the peak of extravascular parasitemia (P1, day 6), and the first trough of the parasitemia wave (T1, day 10). **(C)** Box plot showing quantitative values of vascular permeability corresponding to days 5-8. **(D)** Graphical representation comparing the time of increased *T. brucei* extravasation in each organ (red dot), and the time of increased vascular permeability (blue dot). **(E)** Heat-maps of vascular permeability upon induction by histamine treatment 2 days prior to infection up to day 6 post-infection (left), an extravascular parasite load (right). **(F)** Box plots showing quantitative values corresponding to (F right panel) and the equivalent values in Figure 1G for comparison of extravascular parasite load at days 3-6 post-infection in mice with or without histamine treatment. For all graphs, significance is shown as p < 0.001 (***), p < 0.01 (**), p < 0.05 (*).

The baseline extravascular MFI values in uninfected mice (controls) for FITC-Dextran depends on each organ’s endogenous vascular permeability (**Figure S6A**). g-WAT and isc-BAT were the least permeable organs, followed by the brain, heart and lungs. Spleen, pancreas and lymph nodes showed intermediate levels of basal permeability, while the most permeable organs were the liver and kidneys (**Figure S6B-C**). Organ-specific permeability values in uninfected mice were used to calculate fold changes in permeability throughout infection (**Figure 4**).

Vascular permeability increased in all organs throughout infection (**Figure 4B**). However, we observed significant differences in the extent of such increase and in the day at which leakage was first detected (**Figure 4B-D, Figure S7**). In g-WAT and pancreas, vasculature becomes more permeable from day 5 post-infection, followed by a progressive increase in permeability, and at the end of the infection (day 20), g-WAT is the tissue that shows the highest increase in permeability (14-fold). **Figure 4B** shows organs in increasing order of the onset of increased vascular permeability. Among these organs, the onset of increased vascular permeability happens on day 3 in spleen and lymph nodes, on day 5 in liver, kidney and isc-BAT, on day 6 in brain and heart, and on day 7 in lungs. For most organs, maximum permeability is reached close to the end of the infection (day 20) (**Figure 4B, Figure S7**). Interestingly, in the brain, permeability remains low until day 7, and increases dramatically (2.6 to 5-fold, p < 0.001) on days 8-9, suggesting a significant disruption of the blood-brain barrier around this time. This is consistent with our IVM observations of petechiae and pools of FITC-Dextran across the brain from day 8 of infection onwards (**Figure S7**).

When we compare the variation over time of the increase in vascular permeability of each organ with the extravascular parasite load, we found a significant correlation for most organs (R^2^ values ranging between 0.2 and 0.83; p-values ranging from 0.04 and <0.001), with the exception of the isc-BAT (R^2^ = 0.07, p = 0.24) and lymph nodes (R^2^ = 0.07, p = 0.26). g-WAT and pancreas showed significant correlations throughout the 20 days of infection (R^2^ = 0.42, p = 0.002 for g-WAT; R^2^ = 0.64, p < 0.001 for pancreas) and the highest fold change in the first parasitemia wave (**Figure 4C**). These results suggest that increased vascular permeability favors parasite extravasation in most organs, and that this could contribute to parasite extravasation. However, we noted that the sharp increase in parasite load in g-WAT and pancreas (at day 3 postinfection) (marked in **Figure 4B** by the vertical white dotted line “S” defined in **Figure 1F**) precedes the increase in vascular permeability in these tissues (day 5 post-infection) (**Figure 4D**). Thus, parasites preferentially enter and accumulate in the extravascular spaces of g-WAT and pancreas before vascular integrity is compromised. Considering only days 1-5 of infection, we found no correlation between extravascular parasite load and vascular permeability in g-WAT and pancreas, suggesting that tissue tropism is independent of vascular permeability.

To further confirm that vascular leakage is not involved in the establishment of the preferential reservoirs in g-WAT and pancreas, we induced vascular leakage through pharmacological treatment with histamine, which induces vasodilation and increases leakiness (Egawa et al., 2013) (**Figure 4E left panel**). Upon treatment of mice with histamine 2 days prior to infection with *T. brucei*, and during the first 3 days of infection, the 70kDa FITC-Dextran presence in the organ’s parenchyma revealed a generalized increased vascular permeability, leading to a 1.2 to 5-fold increase in permeability relative to untreated mice during the same days postinfection). We capped the analysis to day 6 post-infection, as this is the first time point by which the tissue reservoirs are already established. In general, the treatment with histamine results in only a very slightly larger number of parasites in the parenchyma of most organs (with only the spleen showing a significant difference relative to untreated mice, p < 0.01) (**Figure 4F**). Importantly, the establishment of the tissue reservoirs in the g-WAT and pancreas starts only on day 3 with a significant increase on day 5, i.e. it does not start earlier than in non-treated conditions (**Figure 4F**).

Altogether we conclude that although vascular permeability is likely a key player for the presence of parasites in the extravascular space of various organs, it does not explain the initial establishment of the reservoirs in g-WAT and pancreas. These results suggest that the mechanism for tissue tropism is active, and independent of vascular permeability.

### Large parasite reservoirs are enriched in endothelial crawlers

*T. brucei* parasites are highly motile and can swim with and against the blood flow or attach to endothelial cells (Bargul et al., 2016; Heddergott et al., 2012; Krüger and Engstler, 2015; Krüger et al., 2018; Uppaluri et al., 2012). Thus, we asked whether differences in intravascular parasite motility could influence or correlate with tissue tropism. To follow how parasites interact with the blood vasculature in different organs, we scored the motility behavior of parasites observed by IVM when blood flow is blocked (**Figure 5**). We found four different types of motility behaviors. The most frequent behavior consists of parasites that mostly preferentially engage with the vessel walls, displaying a constant crawling motility followed by tumbling and crawling in the opposite direction (**Figure 5A top left panel, Supplementary video 3**). Based on previous work (Bargul et al., 2016) we referred to these parasites as ‘crawlers’ (shown in blue in the heat-maps of **Figure 5**). Second, we found parasites that display a long lag time (i.e. no displacement), while persistently probing the endothelial wall with the flagellum (**Figure 5A top right panel, Supplementary videos 4 and 5**). We called parasites displaying this behavior ‘probers’ (shown in red). Then, we found parasites that engage more frequently with RBC than with endothelial cells, showing significantly more tumbling than constant crawling behavior (**Figure 5A bottom left panel, Supplementary video 6**). We called this behavior ‘tumblers’ (shown in black). Finally, we also found parasites that were either barely motile, with only very slight flagellar beating but no productive movement, or fully immotile (shown in grey) (**Figure 5A bottom right panel, Supplementary video 7**). We labeled these parasites ‘immotile’. We hypothesize that parasites that were fully immotile might be dead parasites still not cleared by the immune system.

**Figure 5.**
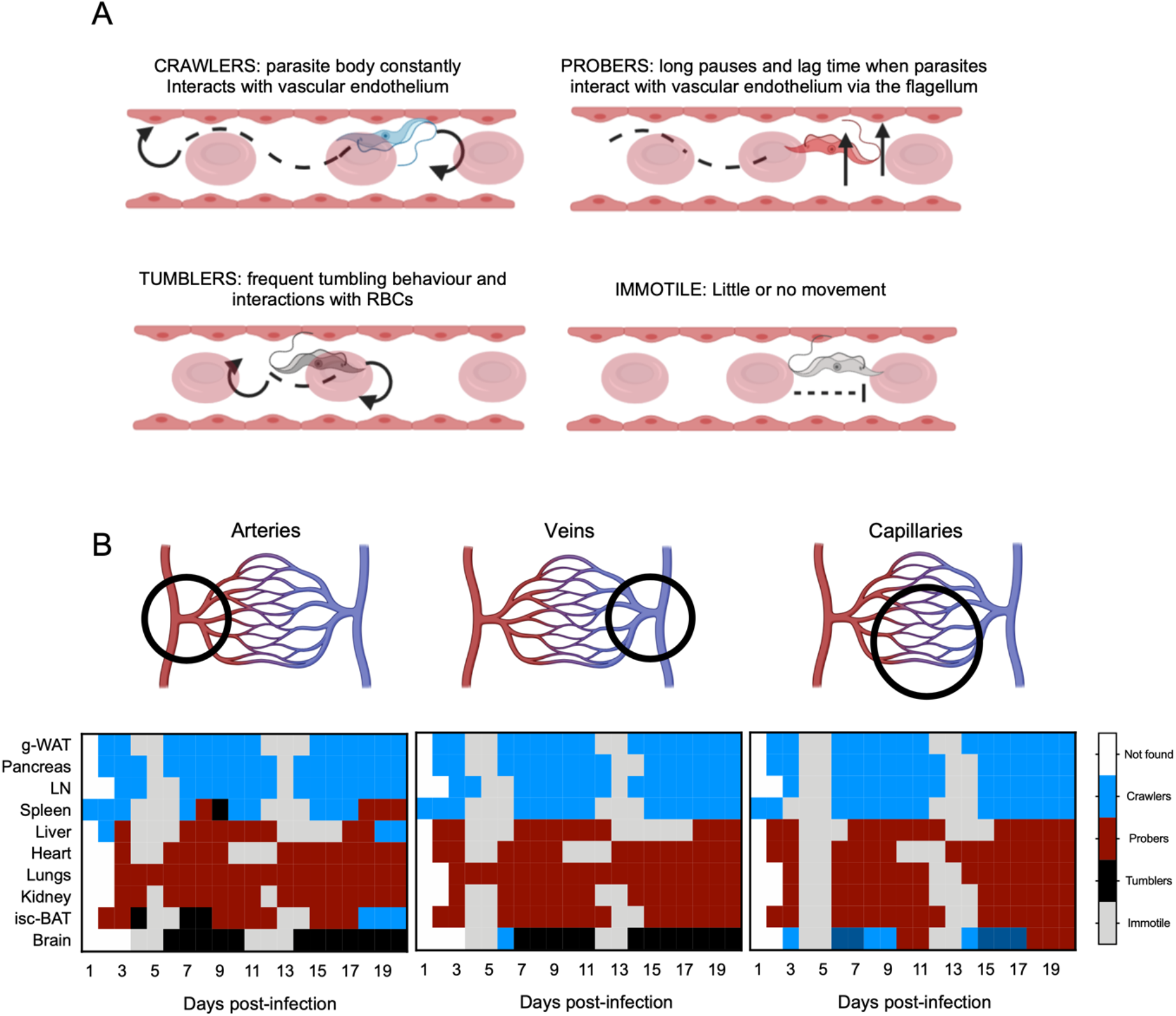
Parasite motility across different organs and vessel types. **(A)** Four types of parasite motility identified *in vivo* upon blocking blood circulation, defined by semiautomatic curation. **(B)** Heat map of dominant parasite motility in each organ, for each type of vessel displayed on the top panel (left-arteries, middle-veins; right-capillaries). Data of frequency of each type of motility in each organ, in each vessel, in each day of infection is shown in Figure S8.

Although we found crawlers, probers tumblers, and immotiles in all organs, the proportion of each phenotype varied significantly according to the organ (**Figure S8**). The dominant type of behavior in each organ (defined as the most frequent population), in each day and in each type vessel type (arterial, venous or capillary) is displayed as color-coded-maps (**Figures 5B**), while the distribution of all motility phenotypes per organ, vascular type and time post-infection is shown in **Figure S8**. Analysis of the dominant phenotype per organ allows us to draw four mains conclusions. First, for most organs, the predominant type of motility remains unchanged throughout infection and is independent of artery/venous origin of the vessel, suggesting that organotypic vasculature has a great impact on parasite motility. Second, two clear groups of organs can be identified: g-WAT, pancreas, lymph nodes and spleen are predominantly populated by crawlers, while liver, heart, lungs, kidney and interscapular BAT are populated by probers. Third, the brain is the only organ in which tumblers are predominant. Remarkably the brain is also the only one evidencing heterogeneity in parasite motility in different vascular beds. Arteries and veins are mainly populated by tumblers, whilst capillaries contain crawlers and probers. Fourth, during the parasite exponential growth and about 1-2 days prior to reaching the first and second peaks of parasitemia (around days 4-5 and 12-13 post-infection), 65-80% of parasites found in the vasculature are immotile (grey shade), which may represent parasites that have already been targeted by the immune system. Outside these two narrow periods, there are no or very few immotile parasites in any organ and any vessel.

Overall, our live imaging analyses show that *T. brucei* parasites display different motility behaviors inside vessels, engaging with both endothelial cells and red blood cells, and that parasites in the g-WAT and pancreas display a similar preferred behavior, namely crawling, although this feature is not unique of these two organs. Nevertheless, given that crawlers were more frequently found in the g-WAT and pancreas, we hypothesize that the frequent interactions with endothelial cells may favour parasite crossing from the intra to the extravascular space of these organs.

### Endothelial adhesion molecules are upregulated during infection

Parasite crawlers establish frequent contacts with endothelial cells, both using the parasite body and the free flagellum. Endothelial cells are covered by many surface molecules that play important communication roles with circulating cells, including leukocyte or cancer cell extravasation (Muller, 2002, 2013; Vestweber, 2015; Wettschureck et al., 2019). Moreover, many of these adhesion molecules have been shown to play a role in host-parasite interactions (Smith et al., 2001). To investigate whether endothelial surface proteins are involved in parasite extravasation, we began by measuring the protein expression levels of six well-known endothelial adhesion molecules in the vessels of all organs: P-selectin, E-selectin (Doré et al., 1993; Hickey et al., 1999), ICAM1, ICAM2 (Lyck and Enzmann, 2015; Rahman and Fazal, 2009), VCAM1 (Muller, 2011; Schlesinger and Bendas, 2015), and CD36. This was achieved by injecting fluorescently labelled antibodies against such adhesion molecules in infected animals at day 6 post-infection, and measuring the fluorescence intensities by intravital microscopy (**Figure 6, Figure S9**). Representative images are shown for g-WAT and pancreas (**Figure 6A**), while other organs are shown in **Figure S9**. The heat-map shows the intensity of antibody staining in animals infected for 6 days relative to non-infected animals (**Figure 6B**).

**Figure 6.**
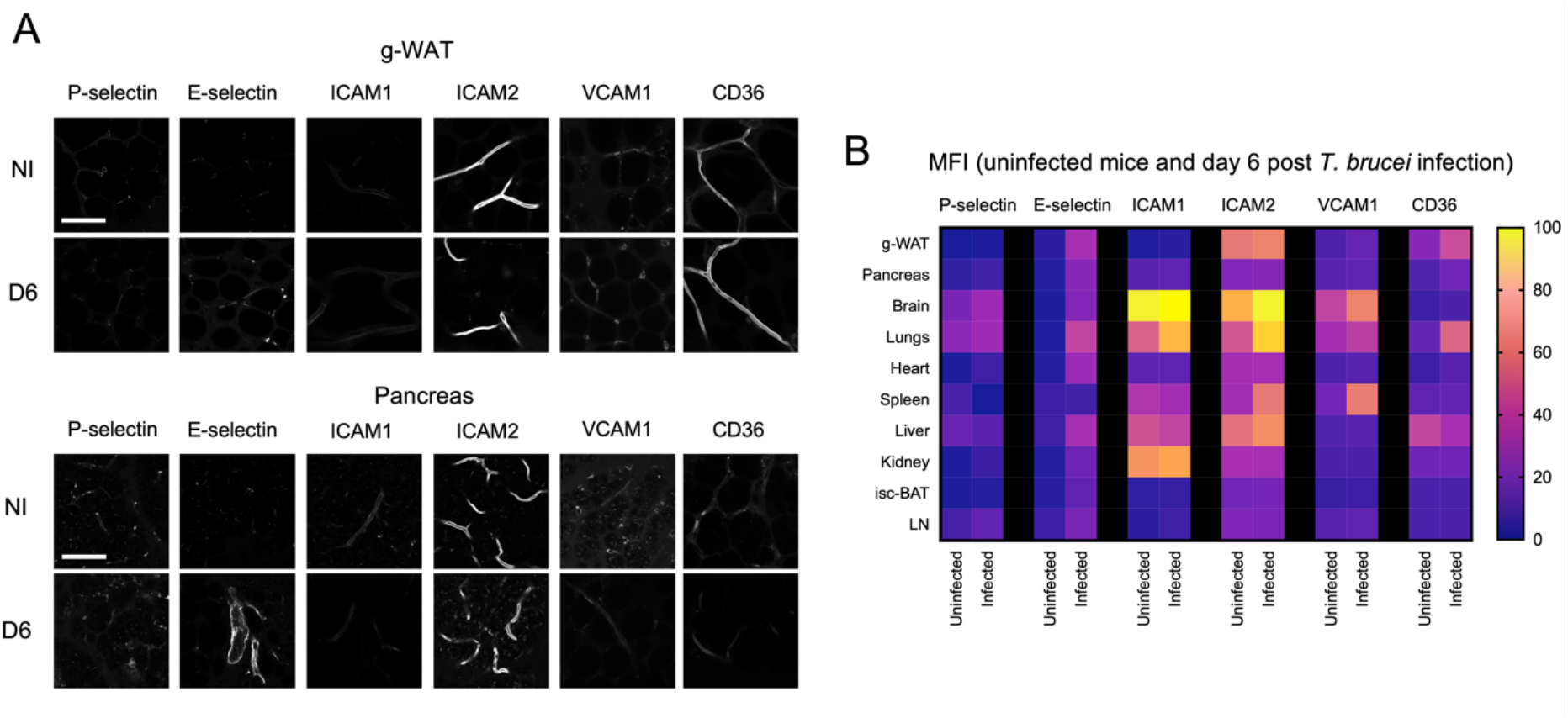
Changes in expression of vascular endothelial receptors during a *T. brucei* infection. **(A)** Expression of 6 endothelial receptors (P-selectin, E-selectin, ICAM1, ICAM2, VCAM1, and CD36) measured by intravital imaging after injection of antibodies coupled to A647 fluorophore. Representative intravital images for (A) relative to g-WAT and pancreas. Scale bar is 50 μm. **(B)** Heat map shows MFI values for each receptor measured in each organ on uninfected and day 6-infected mice..

We observed that several adhesion molecules become more abundant during infection, but the extent of this effect was very variable among organs and among adhesion molecules. Of note, E-selectin is the adhesion molecule that is more up-regulated during infection (11.3-fold increase on average (p < 0.001) ranging from 2.5 to 26.7 across organs). In the infected g-WAT, the most abundant adhesion molecules were CD36, ICAM2 and E-selectin. ICAM2 is stably expressed whilst CD36 and E-selectin were significantly upregulated during infection (1.9- and 10.6-fold, (p values of <0.001)). In the pancreas, E-selectin and ICAM2 are most abundant. Eselectin shows the highest upregulation following infection (13.6-fold, p < 0.001), while P-selectin (1.8-fold, p < 0.001), and CD36 (1.7-fold, p < 0.001) upregulation are also significant.

Given the fact that E-selectin, ICAM2 and CD36 are highly expressed or significantly upregulated in g-WAT and pancreas, they are good candidates to interact with parasites as they crawl through the vessel wall in these tissues.

### Adhesive molecules are necessary for *T. brucei* tissue colonization

We then investigated the effects of blocking each of the highly expressed adhesion molecules (E-selectin, ICAM2 and CD36) on parasitemia and mouse survival. In addition, we targeted P-selectin alone and in combination with E-selectin, as compensation mechanisms for both molecules have been reported (Hickey et al., 1999). We treated mice with the respective blocking antibodies from 2 days prior to infection until 6 days post-infection. We monitored peripheral parasitemia (**Figure 7A-B**). In general, the effect of blocking a receptor led to a significant reduction in parasitemia in peripheral blood, and increased survival (**Figure 7C**). Having observed these strong effects in peripheral parasitemia, we then tested the role of these adhesion molecules in parasite extravasation and reservoir establishment. At day 6 post-infection (day 8 post-treatment), animals were subjected to intravital microscopy. Extravascular parasite load was measured by IVM, and data is expressed in a heat-map (**Figure 7D**) as a fold change relative to (0) control (left panel), and as the percentage of values of WT untreated mice (whose value for each organ is taken as baseline, 100%). Values for g-WAT and pancreas are shown in figure **Figure 7D**, while the remaining organs are shown in **Figure S10**.

**Figure 7.**
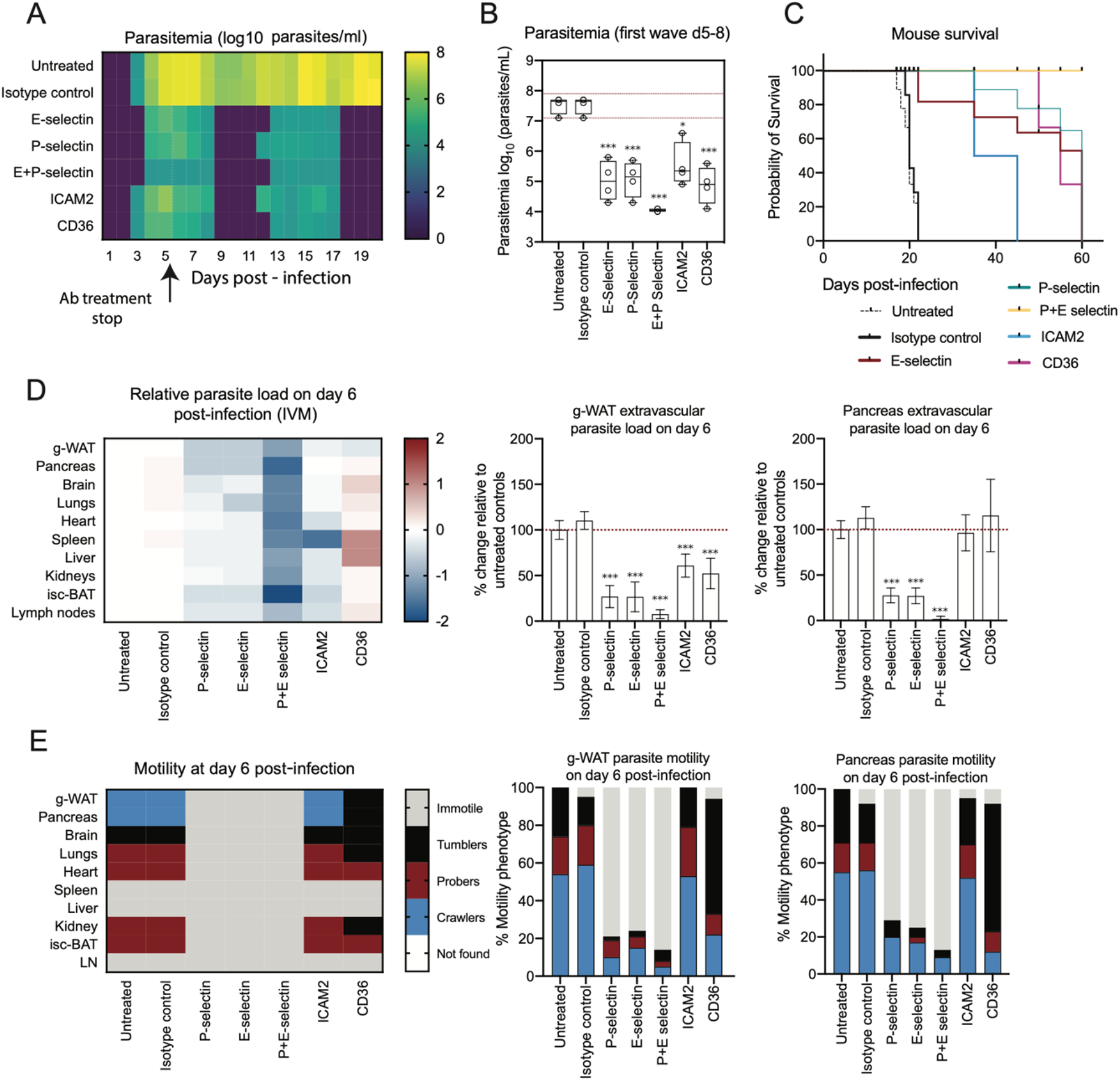
Blocking specific vascular endothelial receptors affects *T. brucei* extravasation. **(A)** Parasitemia was followed by luminescence in mice treated with blocking antibodies 2 days prior to- and during the initial 6 days of infection. **(B)** Box plots showing quantitative values corresponding to (A) during the first parasitemia wave (days 5-8). Significance relative to untreated controls is shown as p < 0.001 (***), p < 0.01 (**), p < 0.05 (*). **(C)** Survival as assessed in antibody-treated mice. Conditions show treatments in which 50% of survival was higher than untreated mice.

Blocking P- and E-selectin resulted in a ~2-fold decrease in parasite load (p< 0.001) in most organs, ~4-fold decrease (p < 0.001) for g-WAT and pancreas; a simultaneous block of both receptors resulted in a synergistic effect with an average drop in parasite load in all organs of ~20-fold (p < 0.001), including g-WAT and pancreas (**Figure 7F**). Consistent with an overall reduction in parasite burden in organs, in the three conditions the peripheral parasitemia was 500-7000-fold lower than controls (p <0.001 for all conditions). In contrast, there was no significant change in parasite load in the pancreas. Next, we measured the effect of this 8-day antibody treatment in the survival of the mice. We observed that blocking E-selectin, P-selectin and ICAM2 allowed 50% of animals to survive past day 40. Interestingly, simultaneous blocking of P- and E-selectin appears to have cured animals since no animal died (**Figure 7C**). Overall, this analysis showed that ICAM2, E- and P-selectins are necessary for parasite extravasation in all organs.

CD36 is the most abundant g-WAT adhesion molecule and its blocking resulted in g-WAT-specific phenotype. While parasite load in the blood decreased 250-fold (**Figure 1B**), it lead to a decrease of 2-fold (p<0.001) in parasite load in the extravascular space of g-WAT, no change in pancreas and a 1.5 to 3-fold (p<0.01) increase in all other organs, including lung and liver where CD36 is also highly expressed (**Figure 7D**). This data shows an organ-specific phenotype, in which CD36 seemingly only affected g-WAT tropism. Consistent with an overall reduction in parasite burden, mice showed delayed lethality (**Figure 7C**). These results strongly support the idea that g-WAT is a major reservoir for *T. brucei* proliferation and/or survival.

Finally, we investigated whether blocking of endothelial adhesion molecules caused any changes in parasite motility inside the vessels that could be associated to a less efficient extravasation. For that, we followed the same procedure described for **Figure 5**, and characterized the most frequent type of motility behavior on day 6 post-infection (**Figure 7E, Figure S11**). When E- and P-selectins were blocked, alone or together, most parasites in all vessels of all organs were mostly immotile, suggesting a rapid clearance of the infection. ICAM2 inhibition did not change parasite mode of motility. In contrast, blocking CD36 affected only g-WAT, pancreas, lungs and kidneys, where the majority of parasite population became tumblers. This data allows us to conclude that parasite motility is highly dependent on the surface proteins that cover the vessel walls in blood circulation, suggesting that parasites engage with such receptors in a cell-to-cell interaction.

Overall, these data show that the organotypic adhesion molecule signature of the host vasculature plays an essential role in parasite fitness and infection outcome. Blocking these endothelial adhesion molecules resulted in a dramatic loss of fitness for the parasite population, as observed by a reduction in parasitemia, and a corresponding reduction in disease burden and mortality for the host. Importantly, CD36 is specifically required for parasite extravasation in the g-WAT, but not in other organs, suggesting that adipose tissue tropism could be mediated by parasite interactions with CD36 at the surface of endothelial cells in adipose tissue vessels.

## Discussion

*T. brucei* are capable of crossing physical barriers separating tissues and the vascular system. Moreover, these parasites have the capacity to preferentially establish extravascular reservoirs in various tissues. By studying the interface between *T. brucei* and the blood vasculature in different organs *in vivo*, we discovered that crossing the vasculature and colonization of solid tissues is necessary for *T. brucei* virulence and greatly affects the disease outcome.

### Parasite reservoirs and pathology

Using a combination of bioluminescence and IVM, we confirmed that the g-WAT is a major parasite reservoir (Trindade et al., 2016), and we identified a previously unknown reservoir: the pancreas. Previous work on intracellular parasites including *T. cruzi* (Corbett et al., 2002; Dufurrena et al., 2017; Martello et al., 2013), *Plasmodium* (Abhilash et al., 2016; Glaharn et al., 2018) and *Toxoplasma* (Nassief Beshay et al., 2018), has reported sequestration in pancreatic blood vessels, pancreatic invasion, and morphological changes in the pancreas, including acute pancreatitis in humans. The fact that the pancreas represents one of the largest reservoirs of an extracellular parasite such as *T. brucei* is puzzling, as the majority of the organ consists of exocrine tissue which produces pancreatic enzymes for digestion including trypsin and chymotrypsin to digest proteins, amylase to digest carbohydrates, and lipase to break down fats (Shi and Liu, 2014). At late times post-infection, we observed organ changes suggestive of pancreatitis, supporting the hypothesis that the presence of the parasite in the pancreas is detrimental for the host. Unlike the white adipose tissue, which vastly reduces in size throughout infection due to lipolysis, the pancreas remains unchanged in terms of size. Moreover, while several major blood vessels supply the pancreas, these are largely common to other organs including the liver, the spleen, and the duodenum, none of which acted as parasite reservoirs, eliminating the possibility that enrichment in the pancreas is simply due to its vascular supply.

Comparative analysis of parasite density in several adipose tissue depots showed that while the g-WAT is the largest reservoir, overall WATs were significantly enriched in parasites compared to brown adipose tissues (BAT). Adipose tissue depots are highly dynamic endocrine organs, with key roles in the regulation of energy metabolism. However brown and white adipose tissues are significantly different in their transcriptional, secretory, morphological and metabolic signatures, and play different roles (Hull and Segall, 1966; Rosell et al., 2014; Saely et al., 2012). We hypothesize that the brown and white adipose tissues present different metabolites to the parasites, the latter of which favour the parasite reservoir. Moreover, in this work, we show that during a *T. brucei* infection, the white and brown adipose tissue depots respond very different in almost all the features we assessed.

In the process of exploring the dynamics of organ colonization we found previously unknown parasite features in each organ. The heart, the brain, the lungs and the kidneys seemed to show high parasite extravasation late in infection, and correlated with vascular pathology. Interestingly, these 4 organs have been associated with specific complications of *T. brucei* infection in humans. Conversely, another group of organs (liver, spleen and lymph nodes) showed high extravascular enrichment early in infection. Importantly these organs are known blood filters, and/or secondary lymphoid organs where immune responses can be initiated and maintained. Research in various pathogens including parasites such as *Plasmodium* and *Schistosoma* have shown early interactions with these organs which ultimately lead to pathology including hepatosplenomegaly and asplenia. Such pathology may also involve organ remodeling including the loss of specific areas necessary for the generation of adequate immune responses (Cadman et al., 2008; Engwerda et al., 2016; Magez et al., 2020; Martin-Jaular et al., 2011; Oster et al., 1980; Urban et al., 2005). In our work, we observed that this early invasion resulted in very high vascular permeability in the liver, and significant enlargement of the spleen and lymph nodes.

### Importance of vasculature in parasite virulence

We showed that neither vascular density nor vascular permeability explain the early colonization of the pancreas and g-WAT. On one hand, both of these organs are relatively less vascularized compared to others showing lower *T. brucei* load. On the other hand, while we noticed significant changes in vascular permeability, these occurred after the establishment of the early reservoirs had already begun (2-3 days earlier). This suggests that the early extravasation of *T. brucei* in these organs is specific, active, and precedes vascular pathology. Given that the intercellular junctions between endothelial cells in arteries are in general much tighter than those in veins (dela Paz and D’Amore, 2009), we hypothesized that *T. brucei*, like leukocytes, might traverse through the path of least resistance (tenertaxis), i.e. the venous vasculature, or the capillaries (Martinelli et al., 2014). It was surprising however, to find that this was not always the case, and that arteries of some organs were enriched in parasites or had significant numbers of extravascular parasites in their vicinity. Specifically in the pancreas, there was a high enrichment irrespective of the vessel calibre, but preferential in vessels of arterial origin, while in the white adipose tissues it was vessels of larger calibre that were enriched. This, as well as the capacity of *T. brucei* to cross the brain blood barrier did not support our hypothesis of tenertaxis, and rather suggests a different mechanism mediating extravasation.

Analysis of parasite motility *in vivo* revealed different swimming phenotypes, consistent with previously described behaviors *in vitro* (Bargul et al., 2016; Krüger et al., 2018) or *in vivo* in zebrafish (Dóró et al., 2019). Importantly, in the pancreas and white adipose tissue, the parasites showed crawling motion along the endothelium, leading to the hypothesis that parasites could interact with adhesion molecules at the surface of the vasculature favoring transmigration. We found that blocking three different surface adhesion molecules in the first week of infection (E- and P-selectin and ICAM2) resulted in a global reduction of parasite load in the blood and solid tissues, which could explain the increased frequency of immotile parasites when selectins were blocked. These results suggest that parasites become more vulnerable to the immune system when interaction with vasculature is blocked. Given the role of these receptors in leukocyte migration, we cannot exclude an alternative explanation, in which the treatment with blocking antibodies might have prevented leukocytes from leaving circulation, which would have resulted in a stronger immune response in the blood. Importantly, the consequence of the 8-day antibody treatment was that animals survived much longer and some even appear to have been cured. These results show that the parasite-vasculature interaction is determinant for parasite virulence. Moreover, it demonstrates that tissue colonization plays an important role in disease severity.

While treatment against E- and P-selectin, and ICAM2 had a global effect on parasite traversal, intravital imaging showed that blocking CD36 resulted in a 50% reduction in *T. brucei* load in the extravascular space of the g-WAT, but no change in parasite load in pancreas, heart, kidneys, and isc-BAT or an increased load in brain, lungs, spleen, liver and lymph nodes. These results show that CD36 specifically favors parasite extravasation in adipose-tissue, which could explain the tropism to this tissue. Importantly, this data shows that the adipose-tissue specific effect of CD36 has a major impact in disease progression, showing for the first time that adipose tissue tropism is a key virulence mechanism.

Importantly, we found that *T. brucei* motility was affected when CD36 was blocked, but not when E- and P-selectin or ICAM2 were blocked. Without access to CD36 in the vascular endothelium of the pancreas and adipose tissues (and other organs), parasite motility switched from crawlers (whereby parasite body constantly interacts with vascular endothelium) to tumblers (frequent tumbling behavior and interactions with RBCs). The loss of the crawling phenotype in CD36-treated mice suggest that this adhesion molecule is important for parasite engagement with vasculature. Meanwhile, in the spleen, liver and lymph nodes, the majority of parasites were immotile and in larger amounts than in a normal infection, suggesting these organs could play a role in eliminating parasites that could not extravasate. Abrogation of CD36-mediated sequestration of *Plasmodium*-infected hosts also results in increased parasite loads in the spleen, as they are more readily eliminated by this lymphoid organ (Fonager et al., 2012; De Niz et al., 2016).

We propose a model for adipose tissue tropism whereby *T. brucei* frequently engages with endothelial surface proteins as parasites travel in the vasculature. Such interactions are not restricted to a specific type of vessel and they can be mediated by several mutually exclusive adhesion molecules (CD36, ICAM2, E- and P-selectins). In the vasculature of the g-WAT, the high expression of CD36 facilitates stronger interactions with *T. brucei*, which swims by frequently interacting with such receptors. It is possible that such interactions may direct or slow down parasites towards a putative crossing site, thus favoring extravasation. Interactions with ICAM2, E- and P-selectin also facilitate extravasation. When CD36 is blocked, *T. brucei* interactions with endothelial cells are reduced, interfering with the typical motility of parasites in the vessels of the g-WAT. This results in less parasite extravasation and more parasites being eliminated from circulation either by systemic immune responses and/or by the spleen.

The fact that parasites and their mammalian hosts have co-existed throughout most of their natural history, has resulted in what is known as a co-evolutionary arms race. This phenomenon promoted the development of parasite mechanisms to evade such responses. Some parasites evolved to be intracellular and use the host cell to disseminate within the host (*Toxoplasma gondii* and *Leishmania* spp.) (Konradt et al., 2016; Peters et al., 2008), others use antigenic variation to periodically change the exposed antigens at their surface (African trypanosomes) or at the surface of infected red blood cells (*Plasmodium* spp.) (Horn, 2014; Rénia and Goh, 2016). A strategy to avoid the destruction by lymphoid organs (Engwerda et al., 2005) is the formation of extravascular reservoirs or sequestration on the vascular endothelium (*Plasmodium* spp.). While the tropism to the skin could favor transmission (Capewell et al., 2016), it remained unclear whether the colonization of other tissues brought any advantage for the parasite. Here we show that tissue colonization is an essential virulence mechanism in *T. brucei.* This is in stark contrast with an evolutionary related parasite, *Trypanosome congolense*, that does not colonize tissue and instead appears to undergo sequestration to vessels (Silva Pereira et al., 2019).

To conclude, in this work unprecedented details on *T. brucei* dynamics *in vivo* in the vasculature of multiple organs, and we demonstrate the key role played by the blood vascular endothelium in the generation and maintenance of tissue reservoirs. The work presented here unequivocally showed that interfering with host vasculature reduced pathology and mortality. This study may pave the way to the development of future novel therapies to tackle this medically relevant neglected tropical disease.

## Acknowledgements

The authors thank Brice Rotureau (Institut Pasteur) for providing the triple reporter (TY1 - TdTomato-FLuc) AnTat1.1^E^ *T. brucei* parasite line. We thank Ruy M. Ribeiro for helpful advice for statistical analysis, and Leonor Pinho for logistical help regarding experimental setups. The authors would like to acknowledge the Rodent Facility, and the Bioimaging Facilities of the Instituto de Medicina Molecular. We thank Marie Ouarne and Maria Angeles Dominguez Cejudo from Claudio Franco’s lab (IMM) for technical support. We also thank Francisca Vasconcelos (Claudio Franco’s lab - IMM), Sara Silva Pereira, Fabien Guegan and Idalio Viegas (Luisa Figueiredo’s lab - IMM) for carefully reading this manuscript and providing helpful and critical feedback. This work was supported by LT000047/2019-L (HFSP) and ALTF 1048-2016 (EMBO) to M.D.N. L.M.F. is an Investigator CEEC of the Fundação para a Ciência e a Tecnologia (CEECIND/03322/2108) and the laboratory is funded by ERC (FatTryp, ref. 771714). C.A.F was supported by European Research Council starting grant (679368), the Fondation LeDucq (17CVD03), the Fundação para a Ciência e a Tecnologia funding (grants: IF/00412/2012; EXPL/BEX-BCM/2258/2013; PRECISE-LISBOA-01-0145-FEDER-016394; PTDC/MED-PAT/31639/2017; PTDC/BIA-CEL/32180/2017; CEECIND/04251/2017).

## Author contributions

M.D.N., C.A.F., and L.M.F. designed the experiments, discussed results, and wrote the paper; M.D.N., D.B., and M.P. conducted the experiments; M.D.N., D.B. and M.P. analysed the data; M.D.N. and A.M.N. established and optimized all protocols relevant to microscopy.

## Materials and Methods

### Animal License and mice used

Animal experiments were performed according to EU regulations and approved by the Animal Ethics Committee of Instituto de Medicina Molecular (AEC_2011_006_LF_TBrucei_IMM). Mice used across this study included wild-type male C57BL/6J mice, and C57BL/6 Albino. All mice were 6—10 weeks old males and were obtained from Charles River, France.

### Mouse infections, parasitemia counts and survival

Mice were infected by intraperitoneal injection of 2,000 *T. brucei* AnTat 1.1^E^ chimeric triple reporter parasites (Calvo-Alvarez et al., 2018) expressing the red-shifted firefly luciferase protein PpyREH9, TdTomato and ty1. For parasite counts, blood samples were taken daily by tail vein puncture, using 1μl of blood diluted in 200 μl of HMI11 medium, and 10 μm loaded in a hemocytometer for parasite quantification. For mouse survival, mice were followed up until clinical signs indicating no recovery. After determining the time of highest disease recrudescence in the survival experiments, all other experiments were capped to 20 days of infection.

### Bioluminescence imaging (*in vivo* and *ex vivo*)

For whole body imaging, infected mice were injected with 200 μl of RediJect D-luciferin (Xenolight, Perkin Elmer) prior to imaging. All measurements were performed in an IVIS Lumina imaging system. A kinetic curve was established to determine the peak of luminescence, which was found to be at 10 minutes post-injection, with a plateau lasting a further 10 minutes. For *in vivo* imaging, mice were anaesthetized with Isofluorane (Isotroy) and imaged using an exposure time of 1 minute and a FOV D (12.5 x 12.5 cm). For *ex vivo* imaging of non-perfused organs, mice were injected with RediJect as described above, and sacrificed using CO_2_ within 3 minutes following this injection. Organs were then extracted, washed in 1x PBS, and placed in a plastic petri dish for imaging at the IVIS Lumina instrument, using an exposure time of 1 minute and a FOV C (10 x 10 cm). For *ex vivo* imaging of perfused organs, mice were injected with RediJect as described above, and sacrificed using CO_2_ within 3 minutes following this injection. The chest was then exposed, the inferior vena cava was cut, and the heart was injected with 40 ml of warm 1x PBS. Organs were then imaged as described above. Image acquisition was obtained, and image analysis performed using the Living Image ® software version 3.0.4.6. For all bioluminescence measures, the results of 9 animals (3 biological replicates in triplicate) are expressed.

### Intravital and *ex vivo* imaging

For intravital imaging, surgeries were separately performed in the brain; the lungs and heart; the liver; the pancreas, spleen and kidney; the adipose tissues and lymph nodes, as described in De Niz *et al* ((De Niz et al., 2019b, 2019c, 2019a, 2020). Briefly, mice were anaesthetized with a mixture of ketamine (120 mg/kg) and xylazine (16 mg/kg) injected intraperitonially. After checking for reflex responses and ensuring none occurred, mice were intraocularly injected with Hoechst 33342 (stock diluted in dH_2_O at 100 mg/ml; injection of 40 μg/kg mouse), 70 kDa FITC-Dextran (stock diluted in 1x PBS at stock concentration of 100 mg/ml; injection of 500 mg/kg), and vascular markers of interest conjugated to A647 (CD31 (Biolegend, used at 20 μg), Ephrin B2 (R&D systems, used at 20 μg), Eph-B4 (R&D systems, used at 20 μg), or VEGFR3 (R&D systems, used at 20 μg). A temporary glass window (Merk rectangular coverglass, 100 mm x 60 mm or circular coverglass (12 mm)) was implanted in each organ, and secured either surgically, with surgical glue, or via a vacuum, in order to enable visualization of the organ surface.

For intravital microscopy, all imaging relative to parasite quantifications, vascular density and vascular leakage was done in a Zeiss Cell Observer SD (spinning disc) confocal microscope (Carl Zeiss Microimaging, equipped with a Yokogawa CSU-X1 confocal scanner, an Evolve 512 EMCCD camera and a Hamamatsu ORCA-flash 4.0 VS camera) or in a 3i Marianas SDC (spinning disc confocal) microscopy (Intelligent Imaging Innovations, equipped with a Yokogawa CSU-X1 confocal scanner and a Photometrics Evolve 512 EMCCD camera). Laser units 405, 488, 561 and 640 were used to image Hoechst in nuclei, extravascular and intravascular FITC-Dextran, TdTomato in *T. brucei*, and CD31, respectively. The objective used to image vascular density, vascular leakage, and proportion of intravascular and extravascular parasites was a 40x LD C-Apochoromat corrected, water immersion objective with 1.1 NA and 0.62 WD. The objective used to classify *T. brucei* movement phenotypes was a 100x plan-apochromat, oil immersion objective with 1.4 NA and 0.17 WD. Between 20 and 100 images were obtained in any one time lapse, with an acquisition rate of 20 frames per second. For vascular density measurements and parasite quantification, in order to gain access to the full organ, we performed *ex vivo* imaging from different organ regions. For this, we performed z-stacks consisting of 16 stacks covering up to 200 μm of tissue depth. For all acquisitions, the software used was ZEN blue edition v.2.6 (for the Zeiss Cell Observed SD) allowing export of images in .czi format, and 3i Slidebook reader v.6.0.22 (for the 3i Marianas SD), allowing export of images in TIFF format.

### Intravascular and extravascular parasite quantification

In order to quantify intravascular and extravascular parasites, we took as reference, the vascular marker CD31-A647 and 70 kDa FITC-Dextran, and we quantified numbers of *T. brucei* parasites within the confines of the regions marked by the vascular marker, and numbers of parasites outside these regions. We then normalized the total quantity to numbers per mm^2^ so as to be able to compare both measurements. Measurements were repeated throughout 20 days of infection, and at least a total of 100 fields of view were quantified.

### Vascular density and diameter quantification

In order to quantify vascular density, we took as reference the vascular marker CD31-A647 and 70 kDa FITC-Dextran. We obtained 100 fields of view, and for each field of view the total area was defined as 100 %. Using the CD31 signal we were able to segment out the vascular regions using Fiji software. We calculated the percentage of vascular area covered using the following formula: 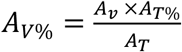, where A_T_ is the total area of the field of view, A_T%_ is 100, and A_TV_ is the total area marked by CD31. Vascular density measurements were performed throughout 20 days of infection. Vessel diameters were measured using Fiji.

### Vascular permeability quantification

In order to quantify vascular permeability changes, we took as reference the marker 70 kDa FITC-Dextran as previously published methodology (Egawa et al., 2013). We measured FITC-Dextran in intravascular and extravascular regions in uninfected mice, and then at each time post-infection with *T. brucei.* The permeability ratio was calculated using the following equation: ***Permeability Ratio*** 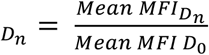, where mean MFI_Dn_ is the extravascular MFI at a specific Day *n*, and the mean MFI_DO_ is the extravascular MFI in uninfected mice (Day 0). To induce vascular hyper-permeability, 5 mg/ml of histamine (Sigma-Aldrich) were prepared in 1 x PBS, and 200 μl were injected intravenously every 2 days, starting 2 days prior to infection.

### Erythrocyte labeling and parasite quantification normalization by vascular type

To quantify and normalize parasite numbers per vascular type, red blood cells were extracted from uninfected mice, and pre-labeled ex-vivo with intracellular dyes CFDA-SE or DDAO-SE as previously described (Theron et al., 2010). The required volume of erythrocytes at 5% hematocrit were resuspended in 500 μl HMI11. RBCs were centrifuged and the pellet resuspended either in 20 μM carboxfluorescein diacetate succinimidyl ester (CFDA-SE) (Invitrogen), or 10 μM 7-hydroxy-9H-(1,3-dichloro-9,9-dimethylacridin-2-one) succinimidyl ester (DDAO-SE) (Invitrogen) in HMI11 and incubated for 20 min at 37°C. The suspension was washed 3x in HMI11, and resuspended at concentrations equaling those observed for parasites at each day post-infection. The labeled RBCs were then injected into infected mice. The number of RBCs per vessel type were quantified, and the number of parasites expressed as a percentage of labeled RBCs in any one vessel as follows:

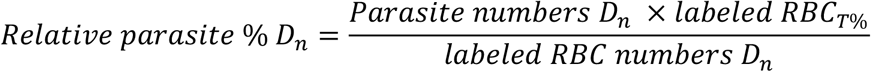

The value of *Relative parasite* % at Day *n* (D_n_) was plotted and color coded in heat maps based on total vasculature, arterial vasculature, or venous vasculature.

### Classification of parasite motility

To visualize parasite dynamic behavior in vessels without flow, we surgically stopped blood flow across different vessels types. We proceeded to image parasites in these vessels, and classified their behavior based on previously established phenotypes (Bargul et al., 2016; Krüger et al., 2018). Parasites were followed for a total of 60 seconds with acquisitions of 20 frames/second, using 100x magnification objectives. Parasites were manually identified as crawlers, tumblers, probers, or immotile. Color-code heat-maps reflect the frequency of each type of parasite motility in the population.

### Labeling endothelial receptors

To investigate the relative expression of endothelial receptors in different organs, we intravenously injected 20 μg of antibodies against E-selectin, P-selectin (BD Pharmingen), ICAM1, ICAM2, CD36 (BioLegend) and VCAM1 (Invitrogen) conjugated to FITC (VCAM1) or A647 (all other antibodies), into uninfected or day 6 infected mice, as previously described in the context of parasitology for CD31 (Hopp et al., 2015). We measured MFIs of at least 100 different vessels per organ in 3 separate mice using an LSM 710 Zeiss microscope, a 40x objective (1.3 NA).

### Blocking endothelial receptors

To investigate the effects of blocking various endothelial receptors, antibodies used included RB40.34 against P-selectin (BD Biosciences, 30 μg per mouse); P2H3 against E-selectin (R&D systems, 20 μg per mouse), a 1:1 combination of both, CBR IC2/2 against ICAM2 (Invitrogen, 20 μg per mouse), and 185-1G2 against CD36 (Abcam, 20 μg per mouse) blocking antibodies were used. Antibodies were injected intravenously daily by tail vein injection, starting two days prior to infection, and continuing until day 6 post-infection.

### Statistical Analysis

Data were displayed in graphs and heat maps generated using Prism 9 software (GraphPad). Means, medians, survival, correlation tests, comparison tests, and error measures were calculated from triplicate experiments with 3 biological replicates each, and/or at least 100 images per condition. For comparisons of bioluminescence measures between organs of different groups we performed multiple t-tests in addition to a one-way ANOVA (differences were considered significant when p < 0.05). Pearson correlations measures (R), and R^2^ values were calculated to determine the strength of linear association between parasite load and either vascular permeability or vascular density. For comparison of proportions between parasites in vessels of different diameters and proportions in vessels of equal diameter across vascular groups, a linear model was performed, and p-values < 0.05 were considered significant. For comparisons of survival, a log-rank (Mantel-Cox) test was performed, and p-values < 0.05 were considered significant. All data used for the generation of the figures is included as a supporting file.

## Supplementary Figure Legends

**Figure S1. Characterization of the triple reporter *T. brucei* line, and parasite load measured by bioluminescence in mice individual organs.**

**(A)** C57BL/6 WT and Albino mice (males, 8 weeks old) were infected with a *T. brucei* reporter line expressing TY1, TdTomato and Firefly luciferase, and their parasitemia (haemocytometer) (left panel), survival (middle panel) and whole body luminescence (left panel) were determined.

**(B)** Bioluminescence imaging of non-perfused organs was performed using an IVIS Lumina system, following injection with RediJect luciferin. While Figure 1 shows the organ mean of 3 separate experiments as a viridis map, Figure S1 shows the mean independent values of each organ separately, as well as the parasitemia curve (in red). Each black line is the log_10_ value of 3 separate experiments with 3 mice each experiment.

**(C)** Schematic showing the location of brown and white adipose tissue depots investigated in this study. Violin plots show the global comparison of parasite loads in WAT and BATs.

**Figure S2. Intravital imaging and *ex vivo* imaging of intravascular and extravascular parasites.**

**(A-D)** Number of parasites per vascular area (normalized to 10 mm^2^) quantified by intravital and *ex vivo* imagining in each organ in the extravascular (blue) and intravascular (red) space throughout infection. Relative parasite distribution in these two compartments is represented in four groups of similar phenotype. **(E)** Schematic representation of groups at early and late times post-infection.

**Figure S3. Changes in vascular density per organ throughout infection.**

**(A)** Vascular density is calculated as the area occupied by the vessel divided by the total area of the field of view were performed following vascular segmentation based on the marker CD31, as shown in Figure 2. This figure shows the individual measurements, with SD for each organ. The red dotted line represents the organ average vascular density throughout infection. **(B)** Representative figures for each organ showing CD31 labeling are included. Scale bar = 50 μm.

**Figure S4. Distribution of *T. brucei* across vasculature of different diameters is organ-specific.**

**(A)** Schematic of vascular branches and respective diameters.

**(B-E left panels)** Heat-maps of parasite distribution in vasculature, relative to labeled RBCs. Organs were grouped by similar parasite distribution.

**(B-E violin plots)** Violin plots showing quantifications of parasite enrichment relative to labeled RBCs (marked as a green dotted line) divided by vessel diameter. For all graphs, significance relative to RBC normalizers is shown as p < 0.001 (***), p < 0.01 (**), p < 0.05 (*).

**Figure S5. Distribution of *T. brucei* across vasculature is organ-specific.**

Organs were grouped by similar parasite distribution.

**(A-D)** Heat-maps of parasite distribution in vasculature, relative to labeled RBCs. Panels are divided as arterial vasculature (left) and venous vasculature (right). Organs were grouped by similar parasite distribution.

**(A-D)** Violin plots showing quantifications of enrichment relative to loaded RBCs (marked as a vertical green dotted line). Plots show distribution of parasites by vessel diameter (0-10, 10-20 and >20 μm) and vessel type (arterial or venous). Data for other organs is shown in Figure 3. Significance between vasculature of different diameters per vessel type is defined as follows: p < 0.001 (***), p < 0.01 (**), p < 0.05 (*).

**Figure S6. Basal vascular permeability in uninfected mice.** Control mice were injected with 70kDa FITC-Dextran (shown in grey values) and Hoechst (blue). **(A)** Intravital images were acquired from 10 organs. Two representative images per organ are included. **(B)** MFI values of the extravascular **(C)** and intravascular niche are included. These values were taken as baseline for all quantifications relative to Figure 4.

**Figure S7. Vascular permeability changes upon *T. brucei* infections.** (**A-H**) Vascular permeability was measured in every organ as an increase in the mean fluorescence intensity (MFI) emitted by the 70kDa FITC-Dextran in the extravascular space. Representative microscopy images of each organ at days 6 and 10 post-infection are shown in the left panels. The rainbow color code reflects fluorescence intensity, with blue being the lowest values and red the highest. The graphics show the vascular permeability of each organ separately through time.

**Figure S8. Distribution of parasite motility phenotypes across organs, vessel types and time of infection.**

Each row represents two organs (divided by a dotted line), while each column represents a vessel type (arterial, venous or capillary). While Figure 5 shows only the dominant motility type per organ and vessel type, Figure S8 shows the distribution of all 4 types of motility.

**Figure S9. Representative images of vascular endothelial receptor expression across organs.**

Each pair of columns represents uninfected and day 6-infected organs., while each row represents a vascular endothelial receptor (P-Selectin, E-selectin, ICAM1, ICAM2, VCAM1, and CD36). Scale bar = 50 μm.

**Figure S10. Relative parasite load in 8 organs following treatment with vascular receptor blocking antibodies.**

Graphs show the respective quantitative values of parasite load in all organs except g-WAT and pancreas (included in Figure 7). Each column represents a condition (blocking antibody), while the red dotted line represents 100%.

**Figure S11. Relative distribution of parasite motility types upon treatment with vascular receptor blocking antibodies.**

Graphs show the respective relative distribution of *T. brucei* motility types in all organs except g-WAT and pancreas (included in Figure 7), upon treatment with blocking antibodies.

## Supplementary Movie Legends

**Video S1. Intravital imaging of *T. brucei* inside mouse vasculature under flow conditions in the g-WAT.** C57BL/6 mouse infected with a *T. brucei* triple reporter line including TdTomato at day 6 post-infection. FITC-Dextran was injected intravenously in the mouse to observe the vasculature and parenchyma (green). *T. brucei* (white) can be observed under flow in the g-WAT. Images were obtained at a rate of 15 frames per second.

**Video S2. Intravital imaging of *T. brucei* inside mouse vasculature under flow conditions in the g-WAT.** C57BL/6 mouse infected with a *T. brucei* triple reporter line including TdTomato at day 3 post-infection. FITC-Dextran was injected intravenously in the mouse to observe the vasculature and parenchyma (green). *T. brucei* (red) can be observed under flow in a g-WAT capillary and in the extravascular space. Images were obtained at a rate of 15 frames per second.

**Video S3. Intravital imaging of *T. brucei* inside mouse g-WAT vasculature with flow stopped (crawlers).** C57BL/6 mouse infected with a *T. brucei* triple reporter line including TdTomato at day 3 post-infection. FITC-Dextran was injected intravenously in the mouse to observe the vasculature and parenchyma (yellow). Multiple *T. brucei* (red) parasites can be observed displaying crawling behavior (image top) while others show tumbling and probing). Images were obtained at a rate of 15 frames per second.

**Video S4. Intravital imaging of *T. brucei* inside mouse g-WAT vasculature with flow stopped (prober).** C57BL/6 mouse infected with a *T. brucei* triple reporter line including TdTomato at day 2 post-infection. FITC-Dextran was injected intravenously in the mouse to observe the vasculature and parenchyma (yellow). One *T. brucei* (red) parasites can be observed displaying probing behavior. Images were obtained at a rate of 15 frames per second.

**Video S5. Intravital imaging of *T. brucei* inside mouse g-WAT vasculature with flow stopped (prober).** C57BL/6 mouse infected with a *T. brucei* triple reporter line including TdTomato at day 2 post-infection. FITC-Dextran was injected intravenously in the mouse to observe the vasculature and parenchyma (yellow). Multiple *T. brucei* (red) parasites can be observed displaying probing behavior. Images were obtained at a rate of 15 frames per second.

**Video S6. Intravital imaging of *T. brucei* inside mouse g-WAT vasculature with flow stopped (tumbler).** C57BL/6 mouse infected with a *T. brucei* triple reporter line including TdTomato at day 3 post-infection. FITC-Dextran was injected intravenously in the mouse to observe the vasculature and parenchyma (yellow). Multiple *T. brucei* (red) parasites can be observed. Parasite at top frame can be observed displaying tumbling behavior. Images were obtained at a rate of 15 frames per second.

**Video S7. Intravital imaging of *T. brucei* inside mouse g-WAT vasculature with flow stopped (immotile).** C57BL/6 mouse infected with a *T. brucei* triple reporter line including TdTomato at day 3 post-infection. FITC-Dextran was injected intravenously in the mouse to observe the vasculature and parenchyma (yellow). One *T. brucei* (red) parasite can be observed immotile blocking a capillary. Images were obtained at a rate of 15 frames per second.

