## Supplementary figures and images for "Organotypic endothelial adhesion molecules are key for *Trypanosoma brucei* tropism and virulence"

### Figure S1

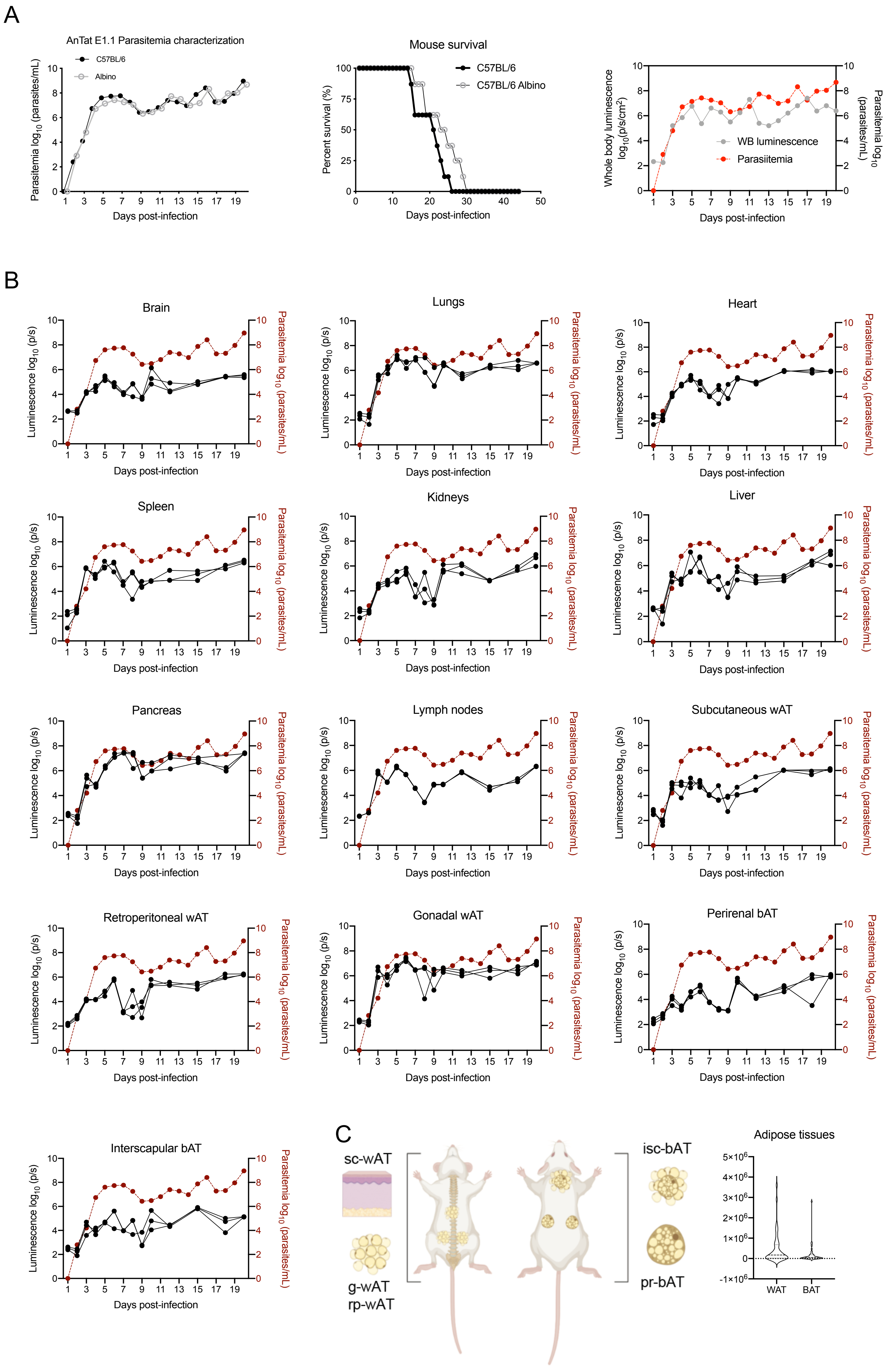

### Figure S2

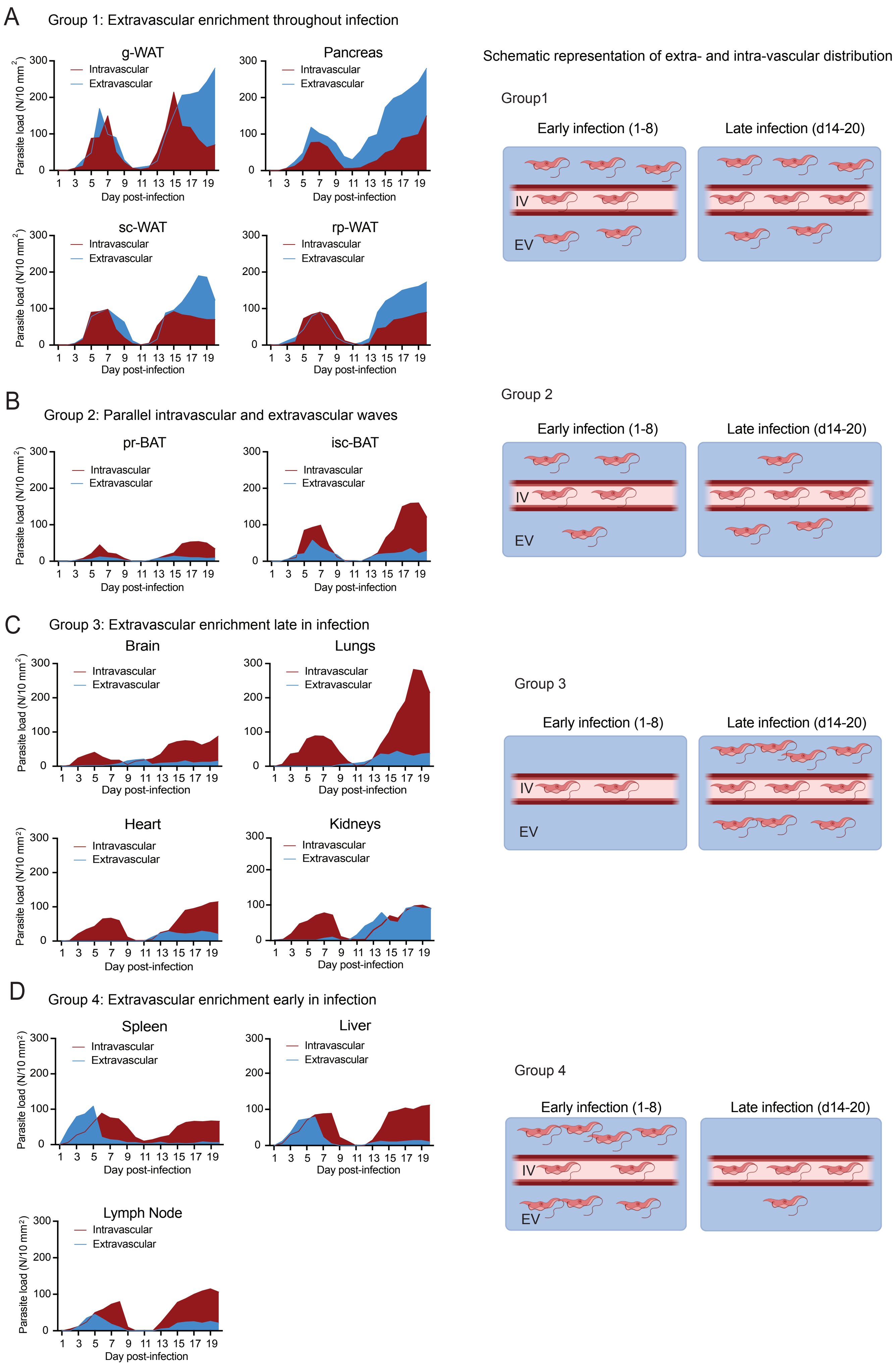

### Figure S3

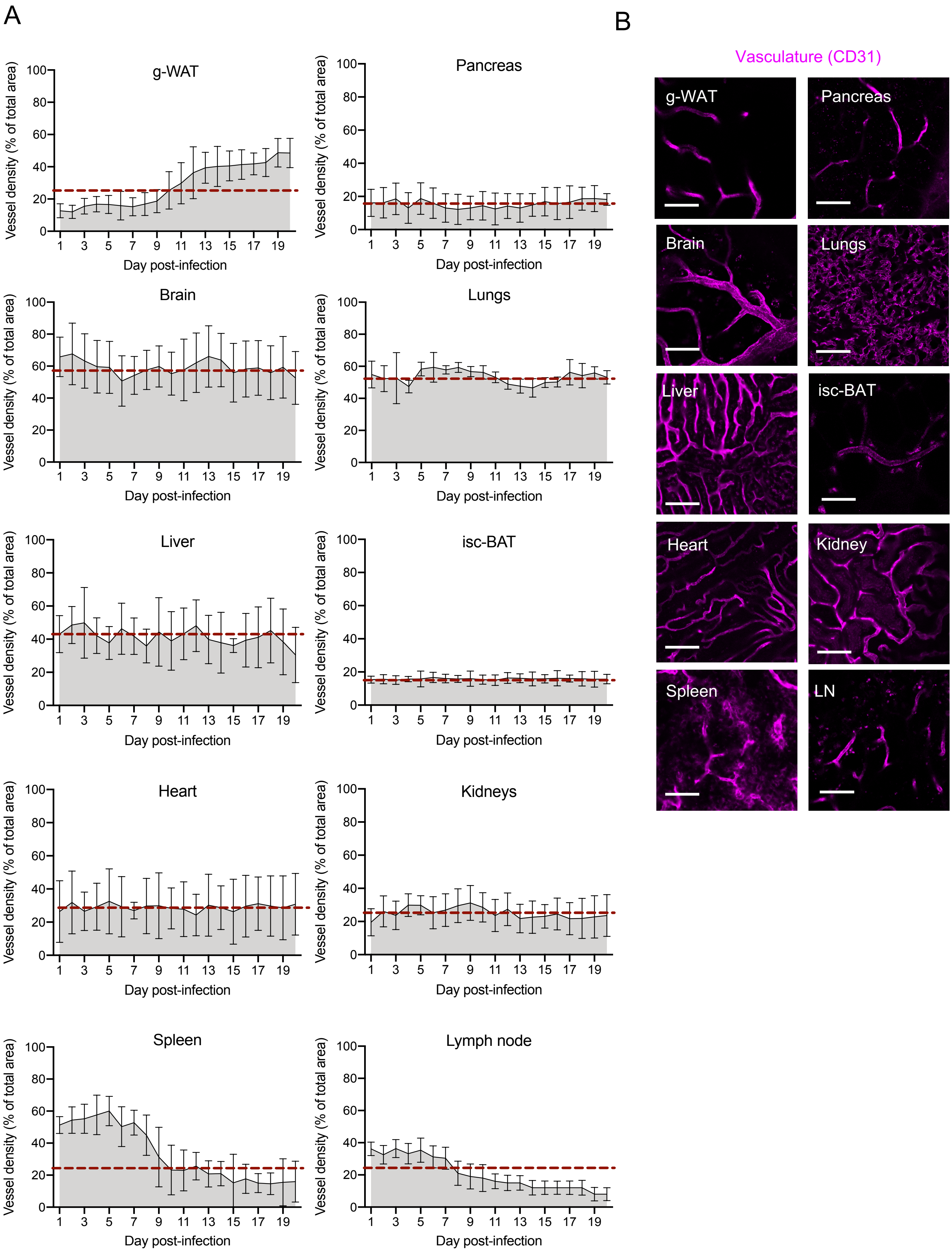

### Figure S4

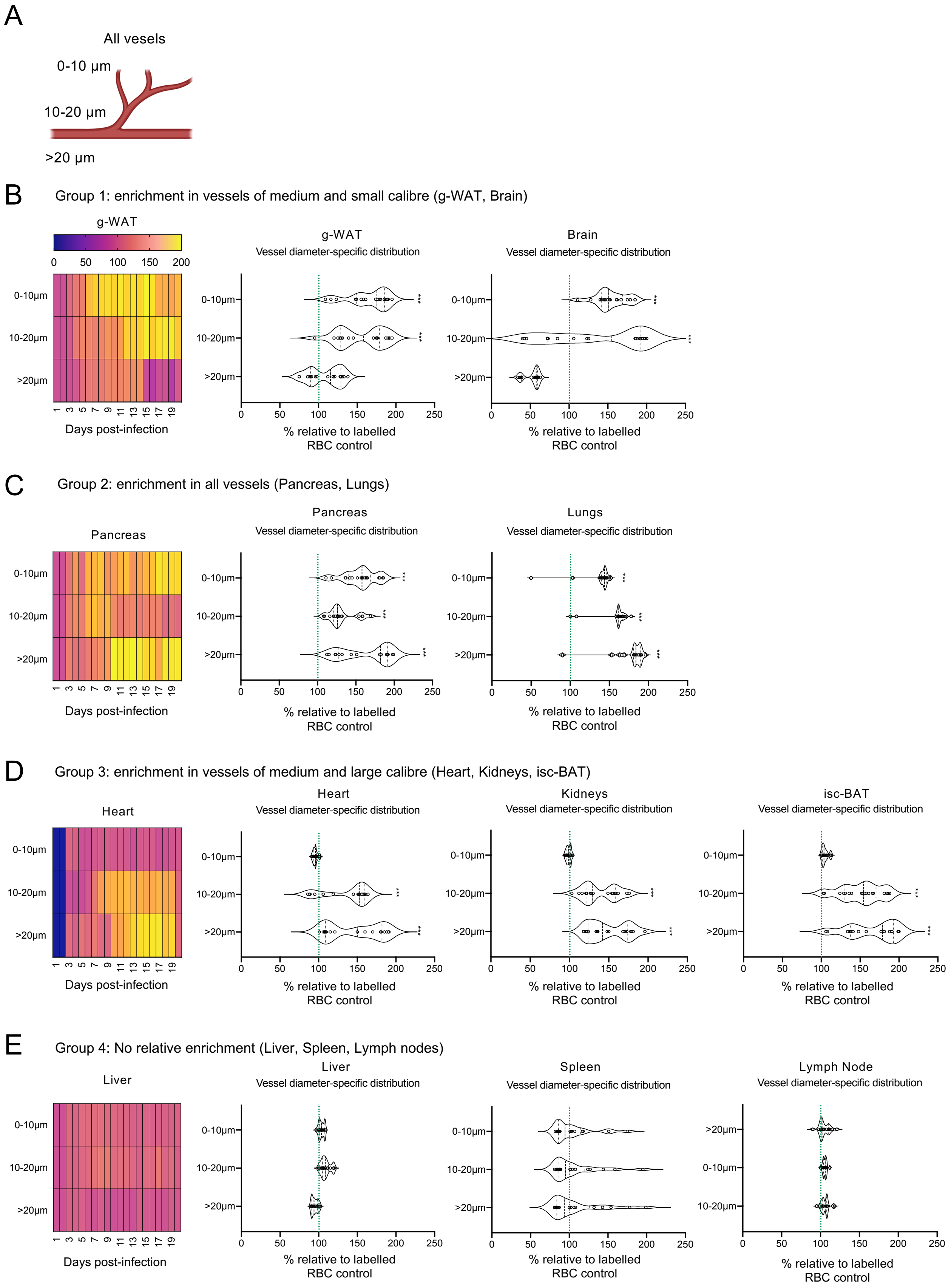

### Figure S5

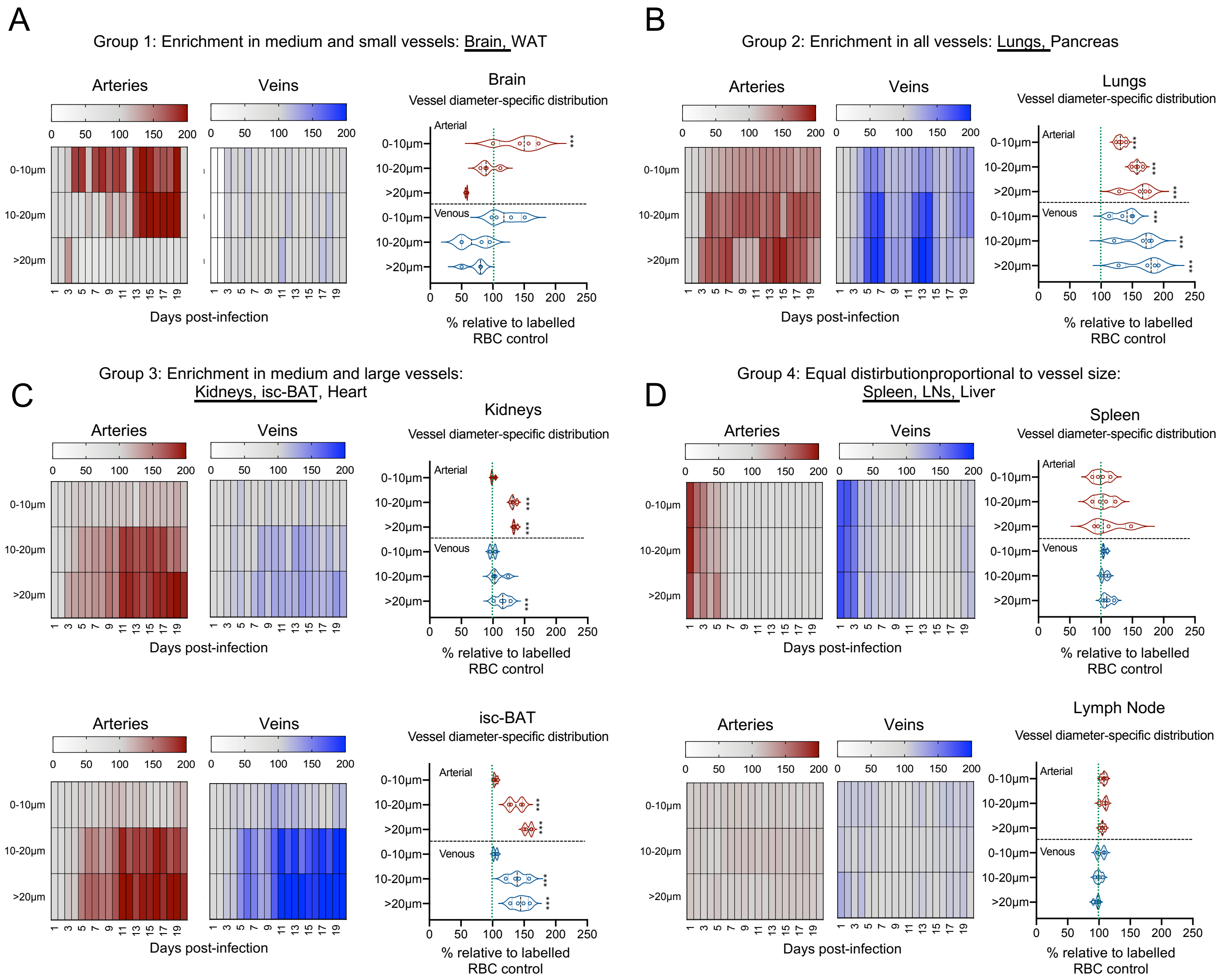

### Figure S6

Figure S6

A

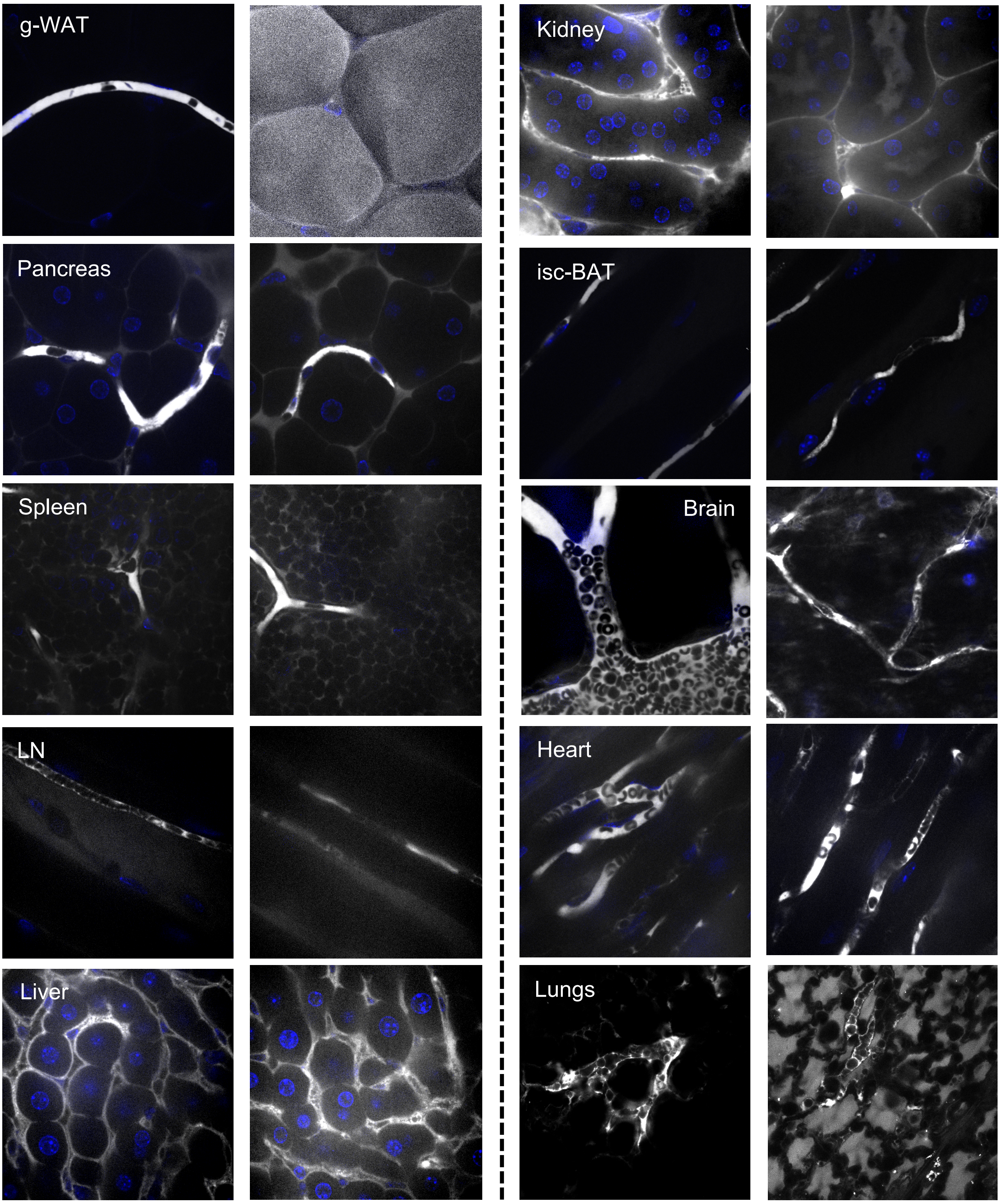


B

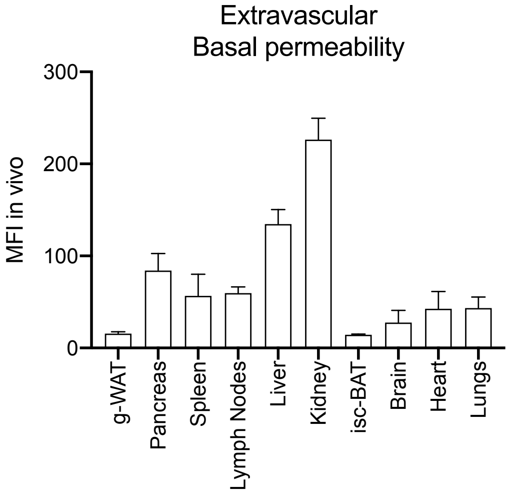


C

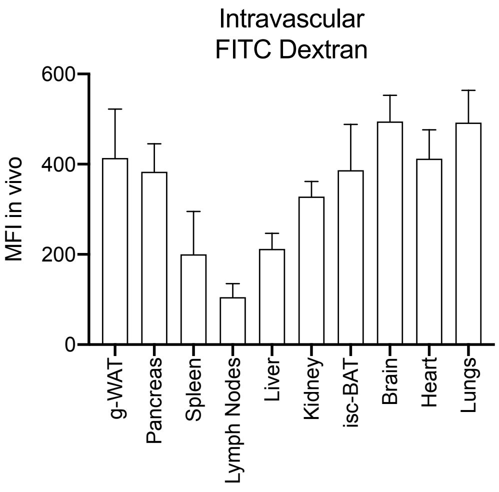

### Figure S7

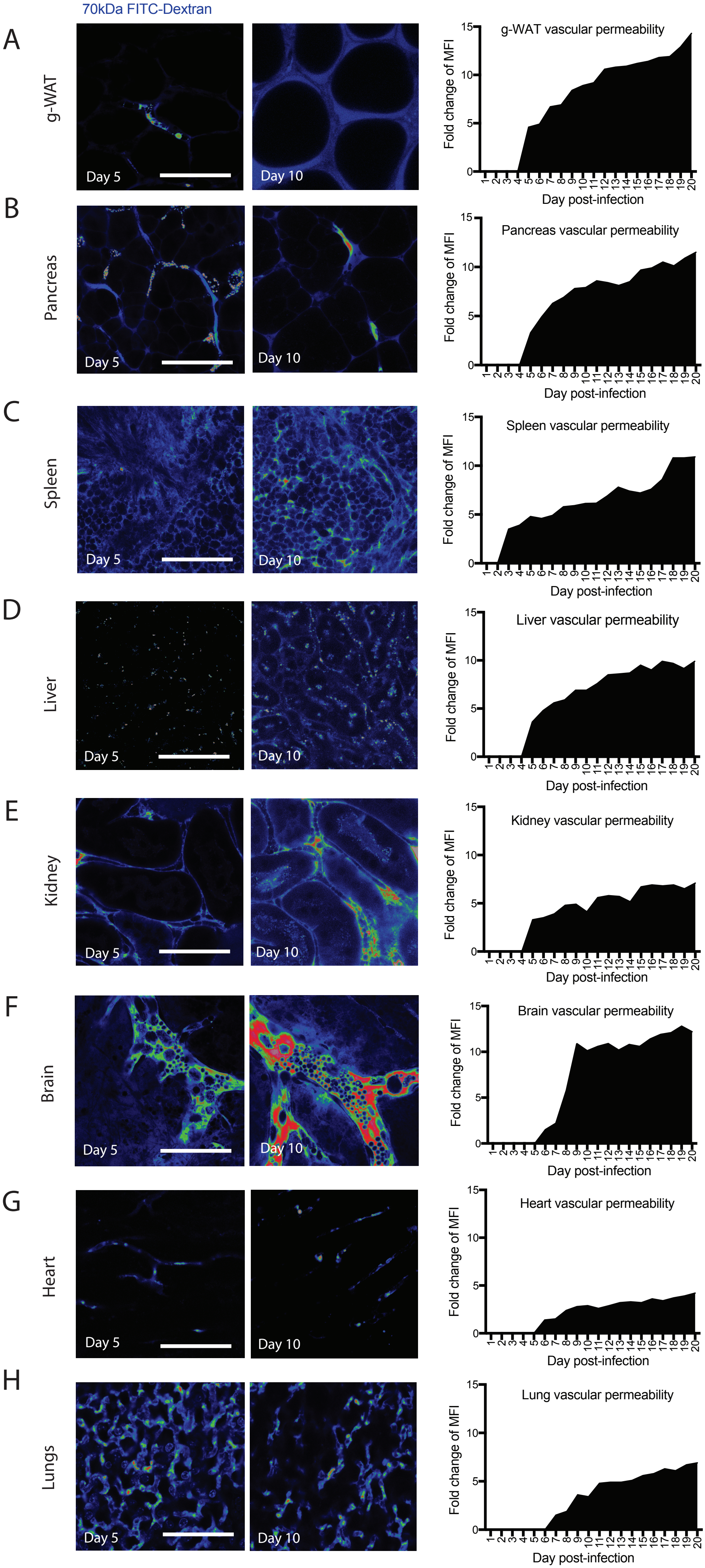

### Figure S8

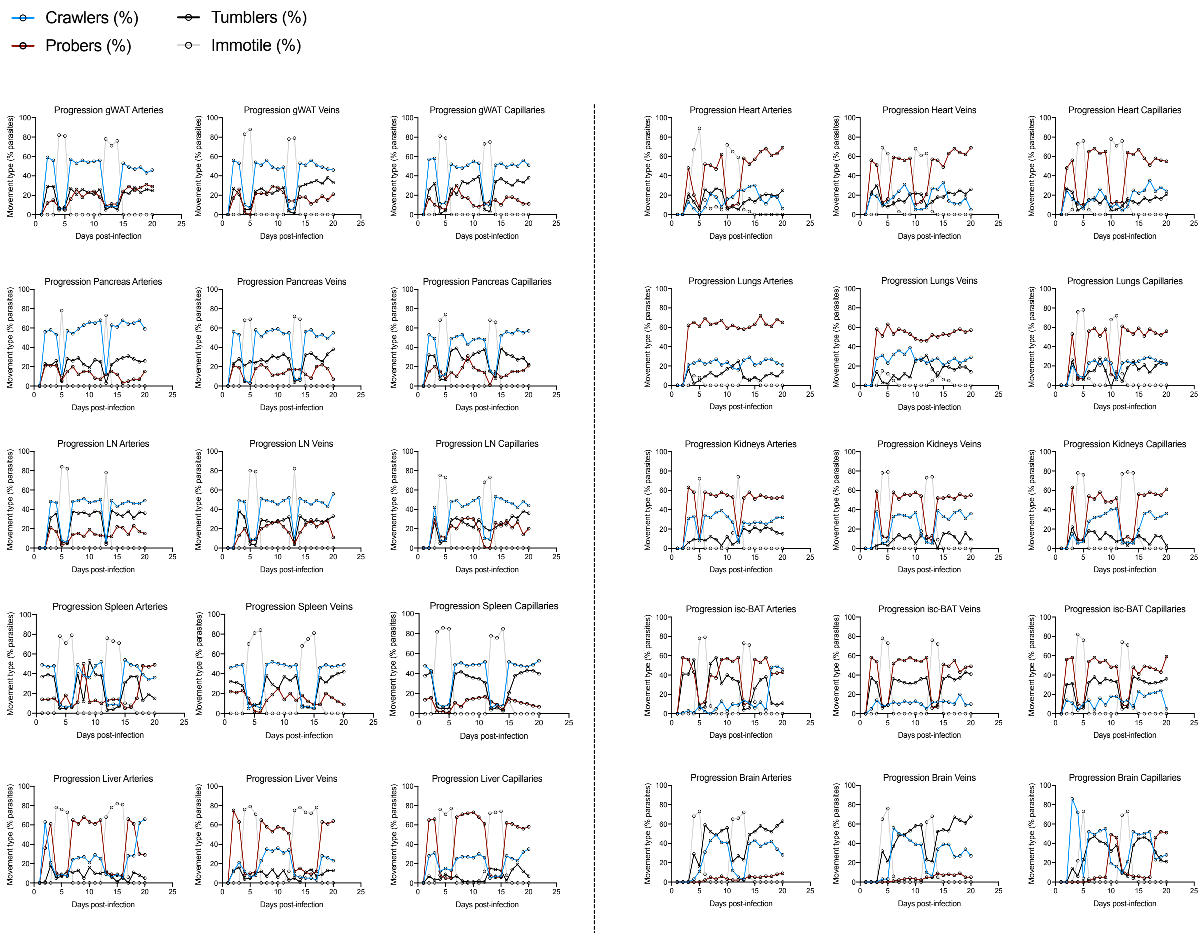

### Figure S9

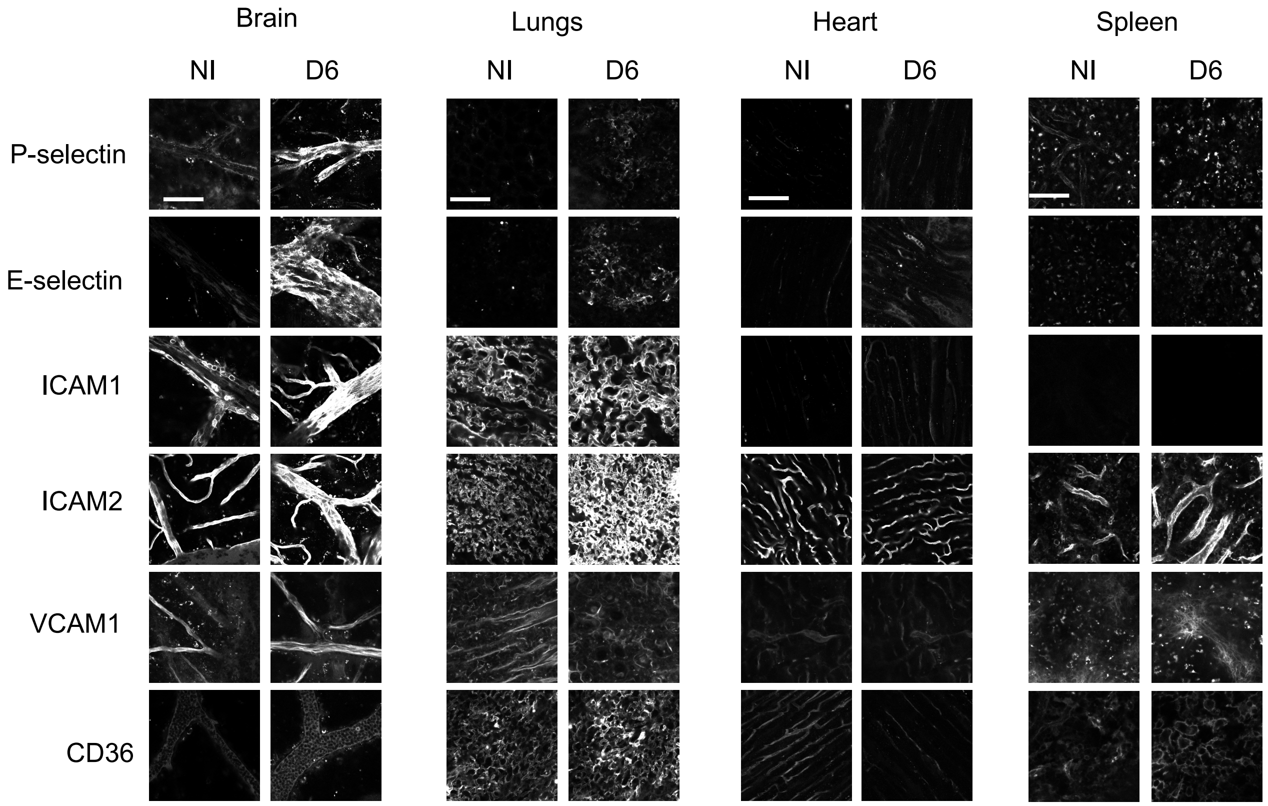

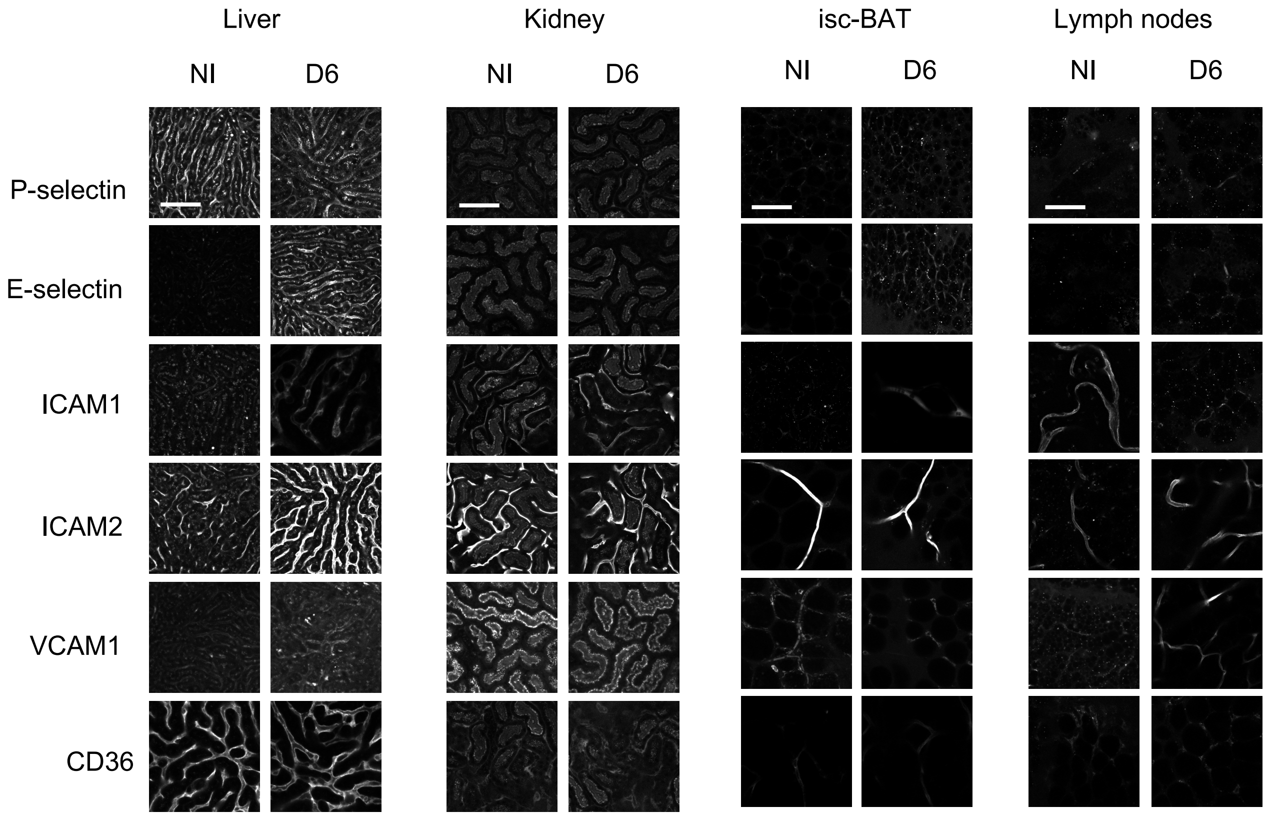

### Figure S10

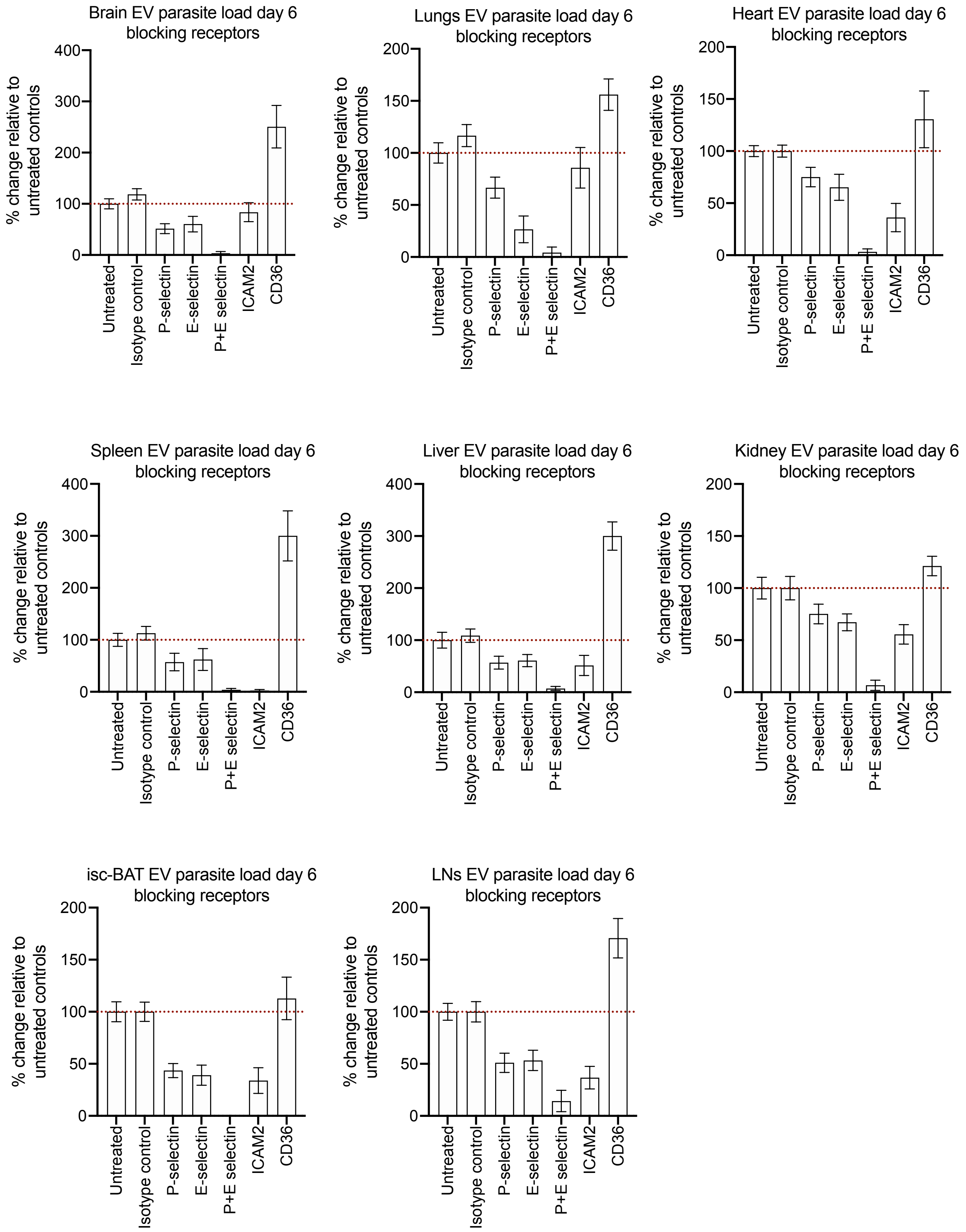

### Figure S11

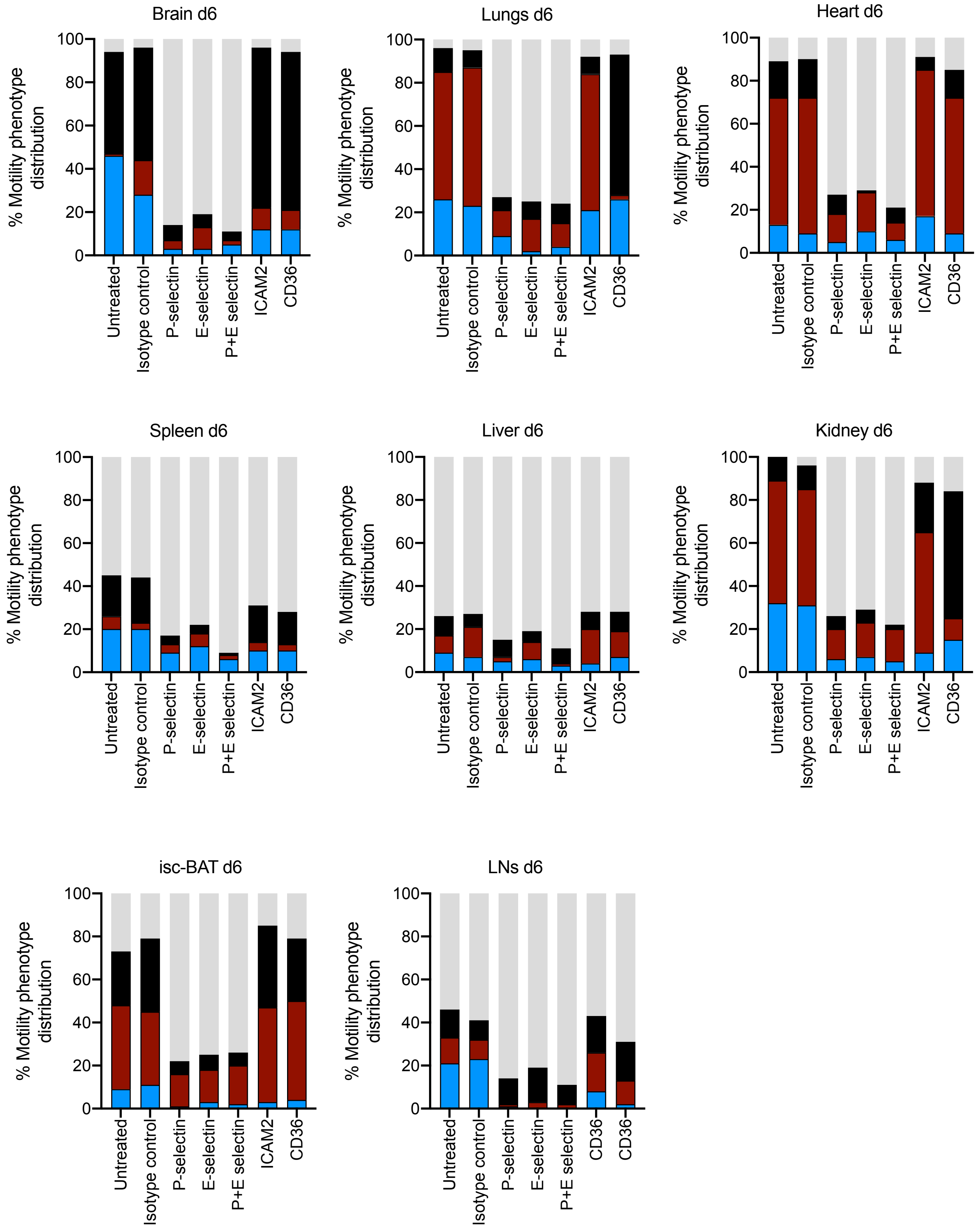
